# Retinal ganglion cell-specific genetic regulation in primary open angle glaucoma

**DOI:** 10.1101/2021.07.14.452417

**Authors:** Maciej S. Daniszewski, Anne Senabouth, Helena H. Liang, Xikun Han, Grace E. Lidgerwood, Damián Hernández, Priyadharshini Sivakumaran, Jordan E. Clarke, Shiang Y. Lim, Jarmon G. Lees, Louise Rooney, Lerna Gulluyan, Emmanuelle Souzeau, Stuart L. Graham, Chia-Ling Chan, Uyen Nguyen, Nona Farbehi, Vikkitharan Gnanasambandapillai, Rachael A. McCloy, Linda Clarke, Lisa Kearns, David A Mackey, Jamie E. Craig, Stuart MacGregor, Joseph E. Powell, Alice Pébay, Alex W. Hewitt

## Abstract

To assess the transcriptomic profile of disease-specific cell populations, fibroblasts from patients with primary open-angle glaucoma (POAG) were reprogrammed into induced pluripotent stem cells (iPSCs) before being differentiated into retinal organoids and compared to those from healthy individuals. We performed single-cell RNA-sequencing of a total of 330,569 cells and identified cluster-specific molecular signatures. Comparing the gene expression profile between cases and controls, we identified novel genetic associations for this blinding disease. Expression quantitative trait mapping identified a total of 2,235 significant loci across all cell types, 58 of which are specific to the retinal ganglion cell subpopulations, which ultimately degenerate in POAG. Transcriptome-wide association analysis identified genes at loci previously associated with POAG, and analysis, conditional on disease status, implicated 54 statistically significant retinal ganglion cell-specific expression quantitative trait loci. This work highlights the power of large-scale iPSC studies to uncover context-specific profiles for a genetically complex disease.

## INTRODUCTION

Glaucoma is the leading cause of irreversible blindness worldwide and experts predict it will affect approximately 80 million people by 2040 ^1^. The most common subtype -- primary open-angle glaucoma (POAG) -- is characterized by an open iridotrabecular meshwork angle and progressive degeneration of retinal ganglion cells (RGCs), which culminates in loss of visual field ^2^. Therapeutic options are currently limited; all are directed at lowering intraocular pressure, which has been shown to slow but not fully prevent or reverse visual loss ^3^.

POAG has one of the highest heritabilities of all common and complex diseases ^4,5^, and much work has focussed on dissecting its genetic architecture. The genetic aetiology of POAG is varied: rare genetic variants, *e.g.* myocilin (*MYOC*)^6^ and optineurin (*OPTN*)^7^, cause disease with high penetrance, while common variants have smaller effect sizes. Genome-Wide Association Analyses (GWAS) have identified over 100 independent loci that carry a common risk allele associated with an increased risk of open-angle glaucoma ^8^. Unlike rare variants that largely contribute to disease by changing protein coding, common variants predominantly act via changes in gene regulation ^9^. Mapping expression quantitative trait loci (eQTL) is one of the powerful approaches used to provide functional evidence of the mechanisms of the common genetic variants, allowing the allelic effect of a variant on disease risk to be linked to changes in gene expression. To avoid spurious associations, and to best understand the cellular effects of changes in gene expression, eQTL mapping needs to be conducted for cells that are pathophysiologically relevant to the disease. In the case of glaucoma, this includes trabecular meshwork cells and RGCs.

The molecular profiling of RGCs in normal and diseased tissue would improve our understanding of the mechanisms that bear a disease risk or contribute to the glaucoma development. Unfortunately, it is impossible to obtain or molecularly profile RGCs from living donors in a non-invasive manner. To overcome this, somatic cells can be reprogrammed into patient-specific induced pluripotent stem cells (iPSCs) ^10,11^, which can then be differentiated into RGCs ^12,13^. Over the years, multiple protocols have been developed to generate RGCs *in vitro* ^14^. Human retinal organoids show a stratified cellular organisation closely resembling the developing human neural retina ^15–20^, and thus it is now possible to generate organoid-derived RGCs for downstream applications, including disease modelling ^13,21,22^ and cell transplantation ^23^. These constructs can also be subjected to single-cell RNA-sequencing (scRNA-seq) to distinguish cell types based on their unique transcriptional signature and examine rare populations that would be missed using bulk RNA-seq ^24–26^. Here, we used scRNA-seq to characterize the transcriptomic profile of the organoid-derived retinal cells, in particular RGCs, generated from a large cohort of the patient-derived iPSCs. We identified a number of cell type and disease-specific eQTLs. Using an additive linear model, a total of 54,786 eQTLs were found to be associated with 21,512 SNPs. By combining our data with recent GWAS results in a transcriptome-wide association study (TWAS), seven genes were identified to be significantly associated with glaucoma development.

## RESULTS

### Large-scale generation of patient iPSCs, differentiation into retinal organoids and scRNA-seq

We recruited a large cohort of individuals, which included healthy (n=57) and patients with advanced POAG (n=55), who were matched by sex. The mean ± SD age at recruitment for controls was : 70.1 ± 8.8 years, and the mean ± SD age at diagnosis for case subjects was 59.7 ± 10.3 years. Participants underwent skin biopsy and their cultured fibroblasts were reprogrammed to iPSCs using episomal vectors as we previously described ^27^. iPSC lines were differentiated to neural retina for 28 days in adherent cultures, after which retinal organoids were then excised, cultured in suspension for 7 days and plated onto Matrigel for an additional 2-week period to allow neuronal outgrowth from RGCs, and harvested for scRNA-seq (**Figure 1A**). Differentiation was performed in batches, with cells harvested and multiplexed for scRNA-Seq with a targeted capture of 16,000 cells from up to eight cell lines per pool.

**Figure 1.**
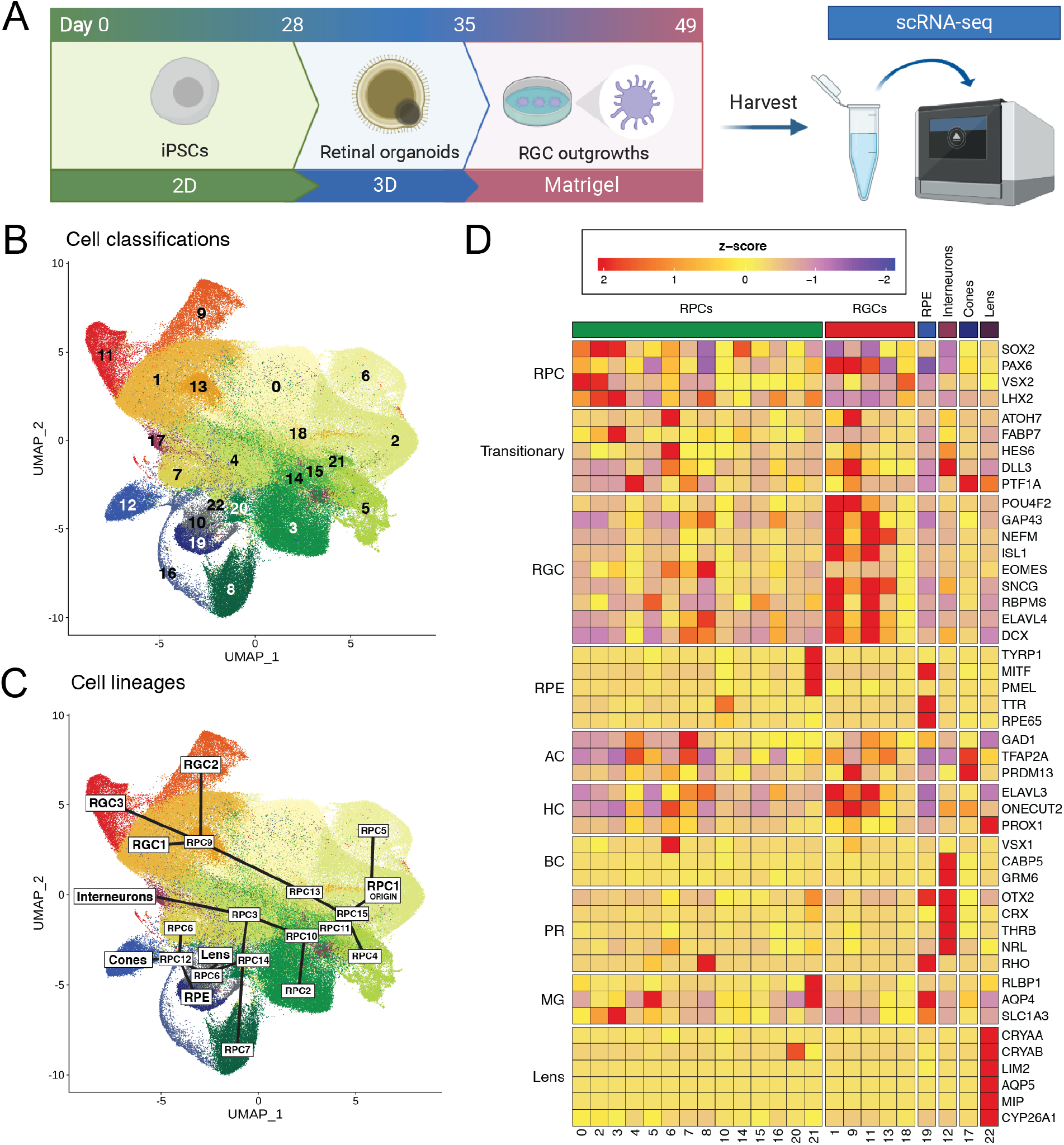
Generation of retinal organoids and identification and characterisation of cell subpopulations. (**A**) Retinal organoids were generated from iPSCs over a period of 49 days. iPSCs were differentiated as a monolayer for the first 28 days, and then cultured in 3D as a suspension for 7 days. Resulting organoids were then plated onto Matrigel and grown until retinal ganglion cells started to project out of the organoid at 49 days. These were harvested for scRNA-seq. (**B**) Uniform Manifold Approximation and Projection (UMAP) representation of cells grouped into 23 subpopulations, as identified by Louvain clustering. (**C**) UMAP plot of the cell types and lineages, as determined by analysis of differentially expressed genes of individual subpopulations and trajectory analysis. RGC clusters form one branch of the trajectory. Other cell types - RPE, interneurons, cones and lens - arise from another branch of the trajectory. The last main branch consists of differentiating RPC subpopulations. RPC: retinal progenitor cell. RGC: retinal ganglion cell. RPE: retinal pigmented epithelium. (**D**) Heatmap of the mean expression of cell type-specific gene markers across in each subpopulation. Expression values have been scaled and converted to z-scores and genes have been grouped by cell type. AC, amacrine cell; BC, bipolar cell; HC, horizontal cell; MG, Müller glia; PR, photoreceptor; RPC, retinal progenitor cell; RPE, retinal pigment epithelium; RGC, retinal ganglion cell.

Twenty-three lines did not differentiate to retinal organoids and were discarded. A total number of 330,569 cells from 160 individual cell lines were captured and sequenced at a mean read depth of 41,020 per cell (**Table S1**). Following cell-specific quality control, 21,346 poor-quality cells were identified and discarded. Genotyping data were generated for all individuals and after quality control and imputation, yielded 7,691,208 autosomal SNPs from 162 individuals at a minor allele frequency (MAF) above 0.01. 514,087 exon-only SNPs were used in conjunction with scRNA-seq data to successfully assign 276,831 cells to 128 donors **(Table S2)**. 32,382 cells were designated doublets or could not be traced to a donor, and were removed from the dataset. Following genotype, virtual karyotyping and scRNA-seq quality control assessment, a total of 24 batches containing 247,520 cells derived from 110 donor iPSC lines were used for subsequent analyses.

### Identification and characterisation of 23 subpopulations from 253,107 cells

Clustering identified 23 subpopulations distributed evenly across cell lines from all donors (**Figure 1B, S1 and S2**). We compared the percentage of cell types between patients with POAG and healthy controls, and observed no statistically significant differences between the groups (p-value ⩾ 0.174, **Figure S2**). Differential expression markers were used to classify the subpopulations to different retinal cell classes based on canonical markers ^28–31^ (**Figure 1C, 1D, Table 1, Table S3**). Retinal progenitor cells (RPCs) represented 77.4% of all cells and localised across 16 subpopulations (**Table 1**). RPCs expressed *PAX6* and *SOX2* transcription factors that are key regulators of neuronal fate ^32,33^, *LHX2*, required for maintenance of open chromatin during retinogenesis ^34^ and gliogenesis ^35^, and the RPC markers *VSX2* and *ASCL1* (**Figure 1D, Figure S1**). Cell cycle genes were not evenly distributed within progenitor subpopulations. The G2/M phase marker *MKi67* was predominantly expressed by cells in RPC1, RPC2 and RPC5. The S phase marker *PCNA* was distributed more broadly; however, the majority of *PCNA*^−^positive cells were identified in RPC clusters 1, 2, 4 and 5 (**Figure 1D**).

**Table 1.**
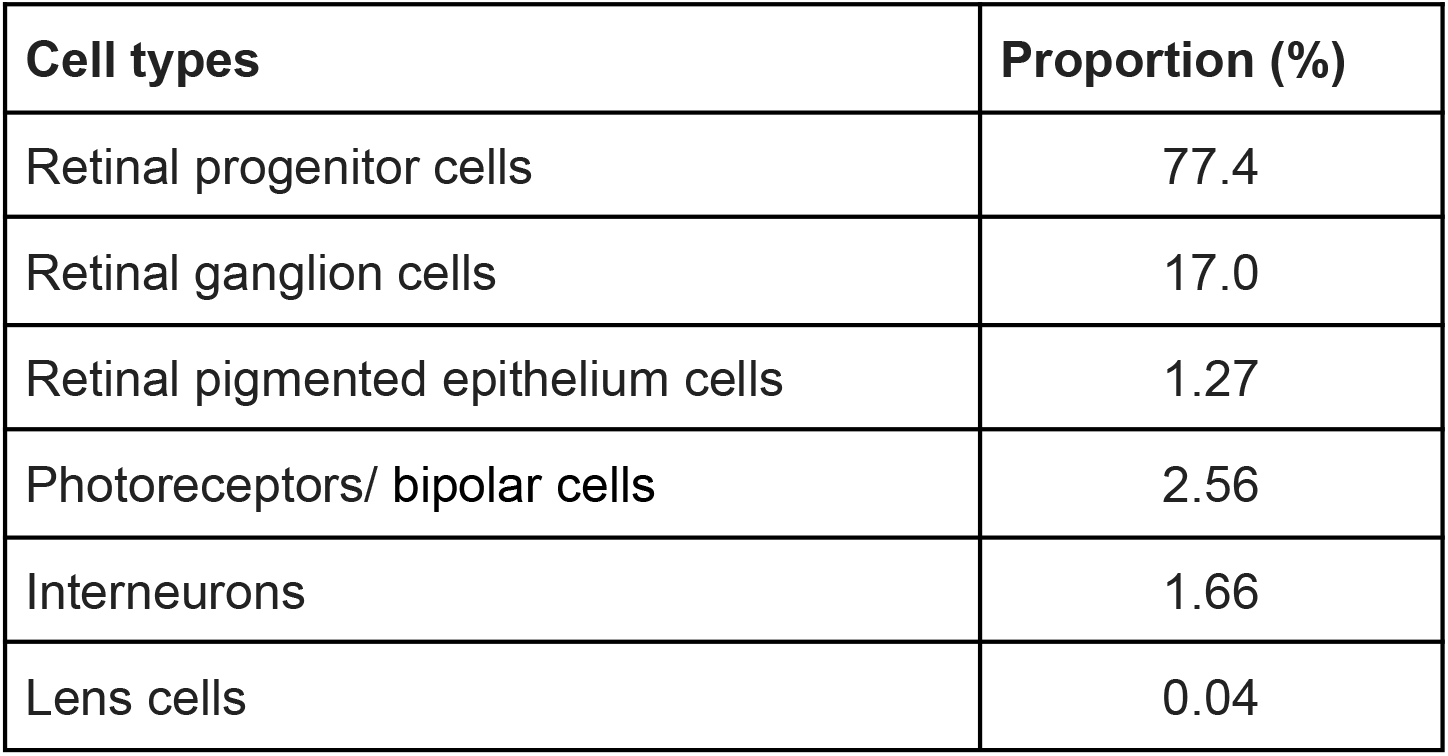
Percentage breakdown of captured cell population. Cells were initially grouped into subpopulations based on Louvain-based clustering. These subpopulations were then annotated using gene markers associated with each known cell type associated with human optic cups.

| Cell types | Proportion (%) |
| --- | --- |
| Retinal progenitor cells | 77.4 |
| Retinal ganglion cells | 17.0 |
| Retinal pigmented epithelium cells | 1.27 |
| Photoreceptors/ bipolar cells | 2.56 |
| Interneurons | 1.66 |
| Lens cells | 0.04 |

RGCs were the second most predominant cell group, representing 17.0% of all cells across the cohort. Based on previous work, RGCs were classified by the expression of the following genes: *POU4F2*, *ISL1*, *RBPMS*, *SNCG*, *GAP43*, *NEFL/M, ELAVL4*, *EOMES* and *DCX ^36–38^*. Three distinct RGC subpopulations (RGC1-3), arising from one subpopulation of RPCs (RPC9), were identified (**Figure 1B-D**, **Table 1**). Pseudotime analysis confirmed the lineage development of RGC1-3 cell types from a single progenitor population (**Figure S3**), with *POU4F2* and *ISL1* showing increasing expression as cells differentiated further from a progenitor state (**Figure 1D**). The expression of both genes are required for RGC specification and differentiation ^39–41^. *POU4F2* expression was generally higher in the RGC1 and RGC2 subpopulations than in RGC3, while *ISL1* expression was higher in RGC1 and RGC3 compared to RGC2. The low levels of *ATOH7* expression in RGC1 and RGC3, in conjunction with the fact that cells in these subpopulations expressed markers of differentiated RGCs such as *SNCG*, *RBPMS*, *GAP43* and *NEFM* (**Figure 1D**), suggests that these subpopulations represent more mature RGCs compared to those from RGC2. We also identified cells expressing markers for photoreceptors/bipolar cells (2.6%) and interneurons (1.7%, **Table 1**). Retinal Pigmented Epithelial (RPE) cells localized in one subpopulation and constituted 1.3% of all cells (**Table 1**). No Müller cells were identified. The various subpopulations are fully described in the **Supplemental Results**. These data are consistent with Sridhar and colleagues ^30^, who also found that RPCs and RGCs are the predominant populations of cells within early retinal organoids.

### The genetic control of gene expression is highly cell type-specific

To explore cell type-specific genetic control of gene expression, we tested for *cis*-eQTL for each cell population independently. We identified a total of 2,235 eQTL across all cell types, which surpassed a study-wide significance threshold of FDR < 0.05 and where the proportion of gene expression in subpopulation-donor pools was > 30% (**Table 2**). The number of eQTL identified per cell type ranged from ten to 456 (**Table S4**), indicating consistent power to detect eQTL and also a similarity of cell types in each population as expected. We assessed the overlap of eQTL between cell types, and found that the majority of cis-eQTL are cell type-specific (**Figure 2A**). Cell-type ubiquitous (shared across cell cell-types) eQTL were rare, with only one eGene (*RPS26*) being common to all cell types. However, we identified a greater degree of shared eQTL amongst related cell types, such as subpopulations belonging to the RGC lineage, specifically RGC1, RGC2, RGC3 and RPC9, which share eQTL for 58 eGenes **(Figure 2B, C).**

**Table 2.** Breakdown of significant cis-eQTLs detected in the full cohort and by disease-status conditional tests. The relationship between genotype and expression was tested at loci within 1MB of each gene, using four different models. Population and disease models were tested using mean expression of all donors, and the population model was used to test control and POAG donors separately. eQTL were labelled significant using the threshold of FDR < 0.05 and gene expression in at least 30% of the donors tested.

| Model | Number of significant eQTLs | Number of significant eGenes | Number of significant eSNPs |
| --- | --- | --- | --- |
| Population | 2,235 | 1,447 | 1,861 |
| Disease | 2,175 | 1,395 | 1,809 |
| Control only population | 1,025 | 804 | 903 |
| POAG only population | 874 | 693 | 770 |

**Figure 2.**
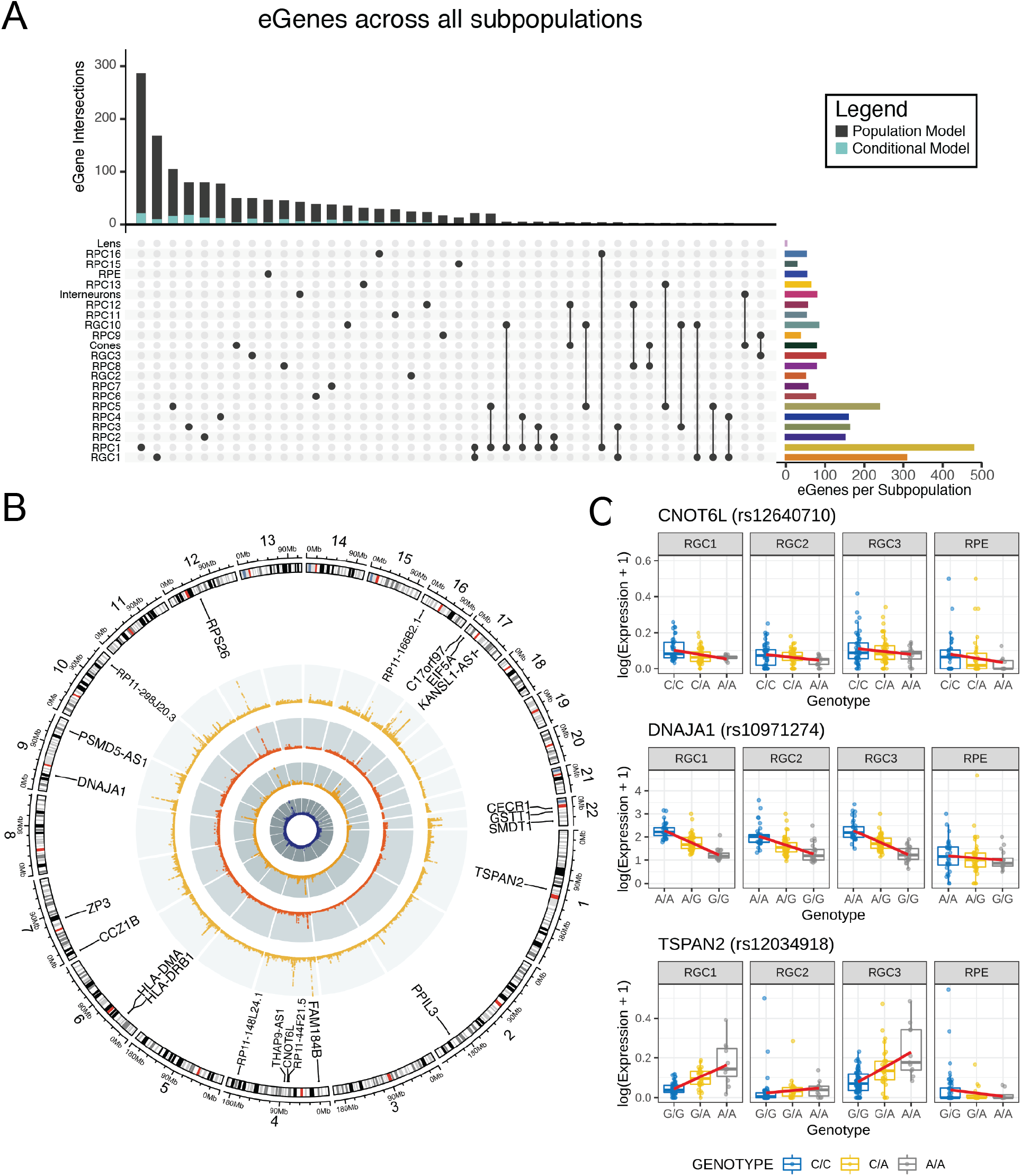
Cell subpopulation-specific eQTLs. Summary of eQTLs specific to cell subpopulations. **(A)** Minimal overlap of genes with significant eQTLs (eGenes) show they are predominantly subpopulation-specific. A small proportion of these eGenes were model-specific. **(B)** Chromosomal map of significant loci in RGC subpopulations RGC1, RGC2, RGC3 and RPE. Loci were labelled as significant if FDR < 5 ✕ 10^−8^. **(C)** Relationships between genotype and mean gene expression at some of the most significant shared eQTL loci in RGC subpopulations - *CNOT6L*, *DNAJA1* and *TSPAN2*, in comparison to corresponding loci within RPE subpopulation.

One potential explanation for the cell type-specific eQTL detection is that the gene is only expressed in one cell type, and therefore, we would not expect to observe an eQTL in the other cell types where the gene is not expressed. To evaluate this, we correlated the expression of each gene that had a significant cell type-specific eQTL effect, with its expression levels in each of the other cell types (**Figure 2C**). These results indicate that cell type-specific eQTL are not a function of cell type-specific gene expression, showing high levels of correlation in almost all instances. Another possible explanation for the cell type-specific eQTL is low statistical power to detect eQTL in multiple cell types. To assess this hypothesis, we implemented an empirical framework to test the rank of the test statistics for eGene SNP effects across the non-significant cell types for each cell type-specific eQTL. In almost all instances we observed none, or very limited, enrichment of the test statistic across cell types.

Among the RGC eQTL identified, a number are directly involved in neurogenesis or neurodegeneration. For example, *CNOT6L* is a poly(A) specific mRNA deadenylase that controls transcription, and reduced expression has been found to impair retinal differentiation in *D. melanogaster* ^42^. *DNAJA1* belongs to a large family of chaperones, and has been shown to prevent neurodegeneration by decreasing α-synuclein aggregates ^43^, whilst *TSPAN2*, is known to support myelination ^44^.

To evaluate the context-specific relationship on the eQTL allelic effect due to disease status, we tested for a SNP-by-disease state interaction and identified 2,175 lead eQTLs (**Table S5**). A total of 290 eQTLs were identified where there was a significant interaction between the SNP allelic effect and POAG disease status. Fifty-four of these eQTLs were specific to RGC lineage subpopulations (**Table S6**). These eQTL are of particular interest, as the data suggest that the allelic effect of the SNP differs due to disease. Interestingly, *rs28368130* at chromosome 9p21, a locus which has been definitively associated with POAG ^8,45^, was found to influence *CDKN2B* expression in the RGC1 cell population with a disease-status FDR p= 4.67×10^−4^. An eQTL for *SPP1* was found across all RGC lineages (**Figure 3A**). *SPP1* encodes for osteopontin, which is known to have reduced expression in eyes from patients with POAG ^46^, and is thought to be neuroprotective to the retina ^47,48^. *IGFBPL1* regulates axonal growth in RGCs ^49^ and was also found to have a statistically significant disease-state interaction eQTL in the RPC9, RGC1 and RGC2 subpopulation of cells (**Figure 3B**). Similarly, *SAR1A*, which is involved in transport between the endoplasmic reticulum to the Golgi apparatus and is associated with axonal growth ^50^, has an eQTL identified by *rs4746023*, in RGC1 cell types. In patients with POAG, carrying each additional copy of the A allele causes an increase by an average of 1.4 transcripts per cell, which is approximately 2 times higher than in healthy controls (**Figure 3C**).

**Figure 3.**
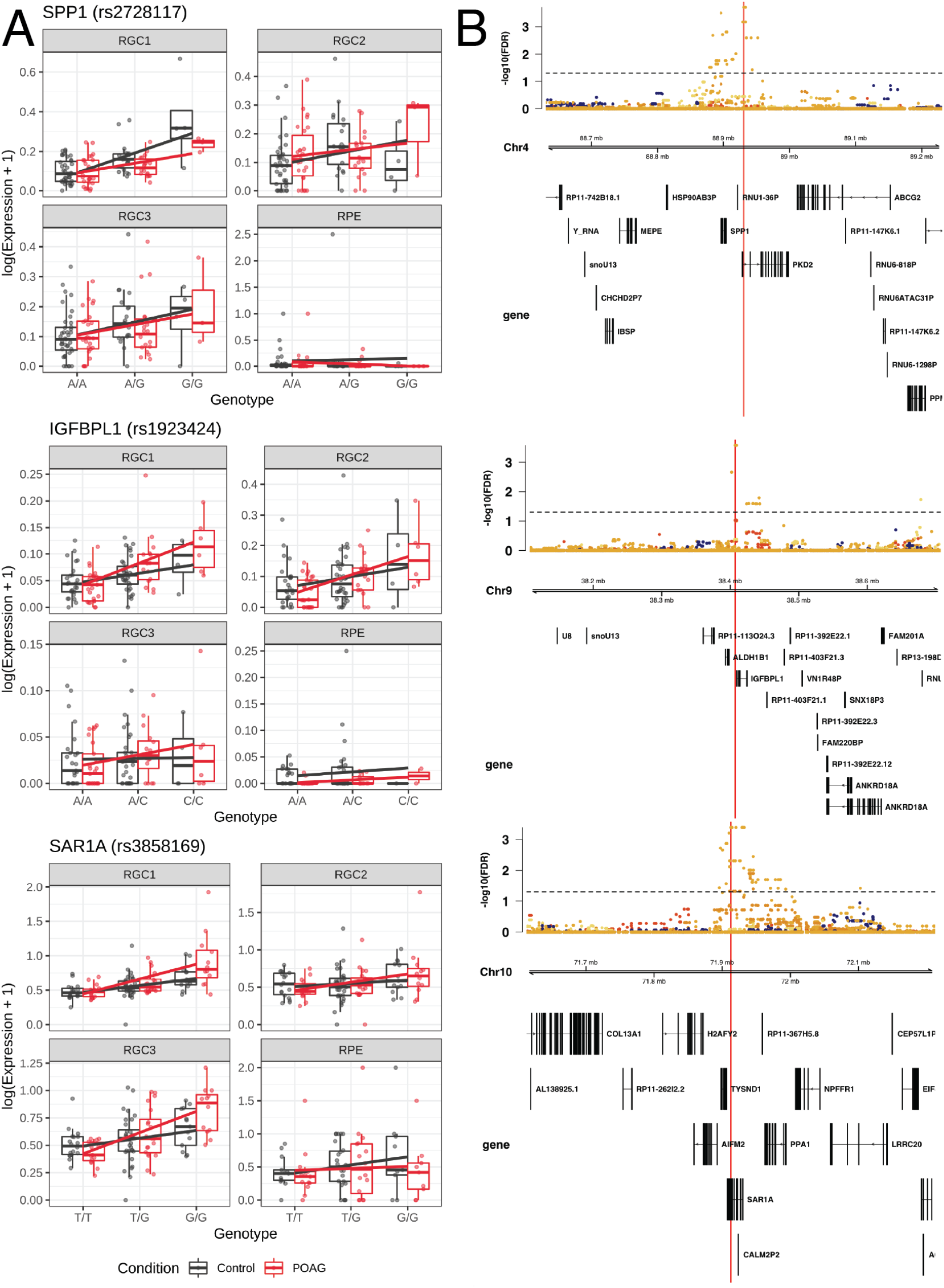
Disease-associated cis-eQTLs specific to cell subpopulations. SNP-by-disease interactions were tested for all SNPs within 1 MBP of each gene for each cell subpopulation. **(A)** The mean gene expression of the following genes: SPP1, IGFBPL1 and SAR1A were grouped by disease and genotype and plotted as boxplots. Regression lines for each condition represent the relationship between genotype and expression. **(B)** Locus zoom plots focus on regions ±30,000bp around locus and genes plotted in the previous panel. FDR values for the significance of eQTL within these regions are plotted by location, alongside the ideograms of genes within this region.

### Disease-specific differential expression of genes across cell types identifies altered transthyretin expression in POAG RGCs

We next sought to evaluate the relationship between disease status and regulation of the transcriptome and a cellular level, testing for differences in the expression levels of genes in each cell population. In total, after Bonferroni correction we identified 3,118 genes whose expression was either up- or down- regulated in POAG cases relative to the controls (**Table S7**). We can be confident that these results are due to the genetic effects underlying POAG risk, as at all steps from iPSC generation, differentiation, cell capture, and library preparation, the cell lines were either managed in either shared conditions, or randomized with respect to disease status (**Methods**). Further, no firm environmental factors have been found to definitively predispose to POAG risk, and are unlikely to account to a difference in gene expression in differentiated cells, given the epigenetic profile of fibroblast-derived iPSCs is reset during reprogramming ^51,52^. Focusing on the three RCG populations, we identified 144 genes differentially expressed between POAG cases and controls (**Figure 4A**). Consistent with our observations of cell type-specific eQTL, 68.06% of genes were identified as differentially expressed in only one cell type, reinforcing the conclusions that the genetic effects of POAG are highly cell type-specific. Interestingly, *TTR* was found to be differentially expressed between POAG cases and healthy controls across all RGC subpopulations (**Figure 4A**). Coding variants in *TTR* are known to cause familial amyloidotic polyneuropathy, which is frequently associated with glaucoma ^53^.

**Figure 4.**
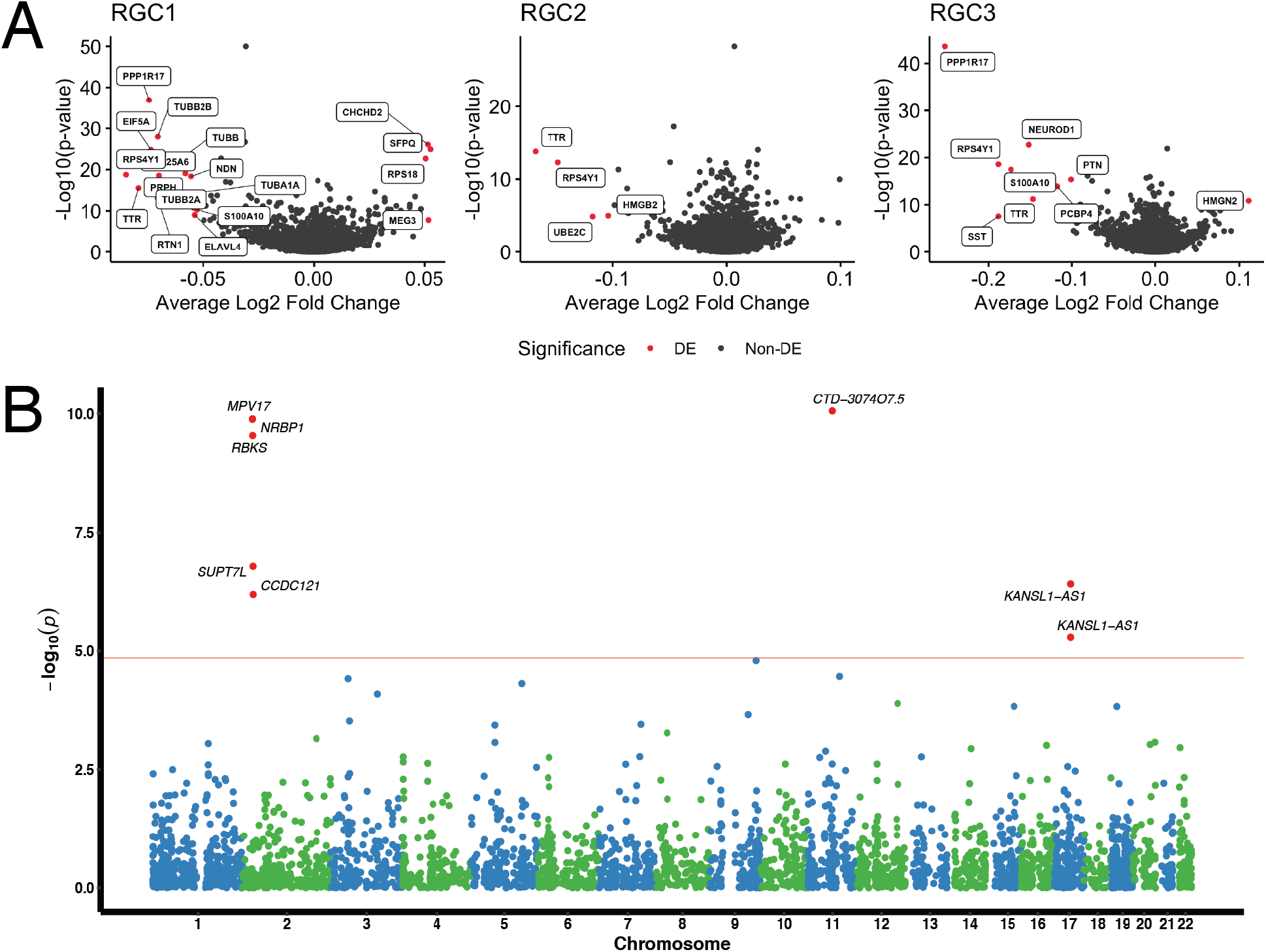
Prioritization of glaucoma risk genes. **(A)** Volcano plot displaying genes differentially expressed between POAG cases and healthy controls (**B**) Manhattan plot displaying the transcriptome-wide association analysis (TWAS) from the scRNA-seq data.

### Transcriptome-wide association study (TWAS) identifies novel and refine known genetic associations for glaucoma

We leveraged this iPSC-derived retinal organoid single cell eQTL data with our recently reported multitrait glaucoma GWAS summary statistics to prioritize glaucoma risk genes in a TWAS ^8^. In the single-cell TWAS, we identified seven genes associated with POAG after Bonferroni correction (**Figure 4B**). Of the five genes identified in the RGC1 subpopulation, one is located at a locus (chromosome 17q21) recently associated with POAG ^54^. Here, we implicate *KANSL1-AS1*, which was also identified as a major eGene for RGCs (**Figure 2B**). *KANSL1-AS1* was also found by TWAS to be associated with POAG in the RPC9 subpopulation.

The TWAS results also helped fine-map potential causal genes at known GWAS loci ^55,56^. Most of the identified genes map to loci that have previously been associated with POAG ^54^. Briefly, the five TWAS genes on chromosome 2 are located near GWAS reported gene *BRE* and share the same GWAS variant rs6741499 (or in strong LD)^8^, which is also associated with IOP (Gao et al., 2018). Of these, the top TWAS hit *MPV17* encodes a mitochondrial inner membrane protein involved in the metabolism of reactive oxygen species ^57^ and has been found to play an important role in the pathogenesis of RGC damage ^58^. The gene *CTD-3074O7.5* on chromosome 11 from RGC2 subpopulation is near *MALAT1*, which is also associated with VCDR ^59^.

We then compared the TWAS results based on scRNA-seq data to bulk RNA-seq data. The bulk retinal transcriptome data were described previously ^60^. Of the three genes with available bulk TWAS results, only *KANSL1-AS1* was significant after Bonferroni correction (P_bulk TWAS_ = 2.95 × 10^−4^).

## DISCUSSION

Here we present a large-scale scRNA-seq analysis of iPSC-derived RGCs. We generated over 100 patient-specific iPSC lines and differentiated them into RGCs using retinal organoids. Following the capture of over 330,000 cells, we analysed 258,071 cells from 112 individuals. We identified a total of 2,235 eQTL across all cell types, and found 58 eGenes that are specific to the RGC lineage. POAG culminates in loss of RGCs, and in iPSC-derived RGCs we identified disease-associated loci. Analysis, conditional on disease status, implicated 54 statistically significant RGC eQTLs, and single cell TWAS identified seven genes at loci previously associated with POAG.

Several recent studies have employed scRNA-seq to characterize transcriptomic changes during human retinal development using fetal tissue ^38,61^, retinal organoids ^62–64^ or both ^30,31^. Our results complement these findings and are in concordance with those of Sridhar *et al.*^30^ who also observed RPCs as the major cell type in early organoids followed by RGCs, photoreceptors and interneurons. Similar observations were made by Lu and colleagues ^31^. The absence of glial cells within the retinal organoids in our dataset is not surprising given these are generally the last retinal cells to develop,^65^ and emerged in older retinal organoids ^30^. Furthermore, we did not observe statistically significant differences in cellular composition of organoids derived from healthy controls and patients with POAG, which suggests a high level of consistency across differentiation batches.

It is becoming widely recognized that the pharmaceutical pipeline for drug development has stalled, and that there is a pressing need for human models of disease to improve our molecular understanding of common, complex diseases and facilitate preclinical trials ^66^. We investigated the impact of genetic background and disease status on gene expression through the eQTL mapping. Highlighting the power of large-scale iPSC studies to uncover disease-specific profiles, this work lays the foundation for context-specific drug screening and underscores the efficiency of using stem cell models for dissecting complex disease.

## Supporting information

Supplemental Results

## ACKNOWLEDGMENTS

We thank Vikrant Singh, Anthony Cook, Alison Conquest and Dominik Kaczorowski for providing technical support. This research was supported by National Health and Medical Research Council Practitioner Fellowships (AWH, JEC, DAM), Senior Research Fellowship (AP, 1154389; SM), Career Development Fellowship (JEP, 1107599) and Investigator Fellowship (JEP, 117578), an Australian Research Council Future Fellowship (AP, FT140100047), an International Postgraduate Research Scholarship & Research Training Program Scholarship (MD), and by grants from the National Health and Medical Research Council (1150144, 1143163) and Australian Research Council (180101405), the Joan and Peter Clemenger Foundation, the Goodridge Foundation, the Ophthalmic Research Institute of Australia, the BrightFocus Foundation, the Philip Neal bequest, Stem Cells Australia – the Australian Research Council Special Research Initiative in Stem Cell Science, the University of Melbourne and Operational Infrastructure Support from the Victorian Government.

## AUTHOR CONTRIBUTIONS

Conceptualization, J.E.P., A.P., A.W.H.; Methodology, M.S.D., A.S., X.H., S.M., J.E.P, A.P., A.W.H.; Investigation, M.S.D., H.H.L., X.H., G.E.L., D.H., L.R., P.S., J.E., CL., U.N., R.M., V.G., S.Y.L., J.L., L.G.; Resources, H.H.L., E.S., S.L.G., L.C., L.K., D.A.M., J.E.C., J.E.P., A.P., A.W.H.; Data analysis, M.S.D., A.S., X.H., S.M., J.E.P, A.P., A.W.H.; Writing - original draft, M.S.D., A.S., X.H., S.M., J.E.P, A.P., A.W.H.; Writing - review & editing, M.S.D., A.S., X.H., G.E.L., D.H., D.A.M., J.E.C., S.M., J.E.P., A.P., A.W.H.; Supervision and project administration, J.E.P, A.P., A.W.H.; Funding Acquisition, D.A.M, J.E.C, S.M., J.E.P., A.P., A.W.H.

## METHODS

### Participant recruitment

All participants gave informed written consent ^78^. All experimental work was approved by the Human Research Ethics committees of the Royal Victorian Eye and Ear Hospital (11/1031H, 13/1151H-004), University of Melbourne (1545394), University of Tasmania (H0014124) and the University of Western Australia (RA/4/1/5255) as per the requirements of the National Health & Medical Research Council of Australia (NHMRC) and in accordance with the Declarations of Helsinki. We recruited a large cohort of patients with POAG and sex-, ethnically, age-matched individuals, through the Glaucoma Inheritance Study in Tasmania and the Australian and New Zealand Registry of Advanced Glaucoma, local ophthalmic clinics and adjunct studies (mean ± SD age: 59.7 ± 10.3 years at diagnosis for case subjects; 70.1 ± 8.8 years at recruitment for controls). To refine translational significance, recruitment of patients with POAG focused on a) end-stage disease, which carries the greatest visual morbidity: virtually the entire optic nerve function is lost through RGC degeneration; b) patients designated as having Normal Tension Glaucoma, as they have RGCs with an increased susceptibility to degeneration compared to cases with trabecular dysfunction or a very high intraocular pressure; c) “super selection” of the most severe POAG cases. Case-inclusion criteria were: in the worst eye a vertical cup:disc ratio >0.95 and a best-corrected visual acuity worse than 6/60 due to POAG or on a reliable Humphrey Visual Field a mean deviation of ≤-22dB; or at least 2 out of 4 central squares involved with a Pattern Standard Deviation of <0.5%. The maximum documented pre-treatment IOP, measured by Goldmann applanation tonometry, was recorded. To fulfil a standard clinical diagnosis of Normal Tension POAG the maximum-recorded IOP was required to be <22 mmHg. Clinical-exclusion criteria were signs of secondary or syndromic glaucoma. For each control participant, a complete ophthalmologic evaluation (incorporating automated visual field testing, fundus and optic disc imaging, corneal pachymetry) was performed. An ophthalmic history was obtained, with questions centred on age at diagnosis, family history, surgical intervention for glaucoma or cataract, macular degeneration, retinal detachment, and refractive surgery. Control subjects, with no known family history of glaucoma, and who had normal IOP, optic discs (optical coherence tomography retinal nerve fibre layer analysis within age-matched normal limits), and visual fields, were selected for analysis.

### Fibroblast culture

Skin biopsies were obtained from non-sun exposed regions using a 3mm^2^ dermal punch. Fibroblasts were expanded, cultured and banked in DMEM with high glucose, 10% fetal bovine serum (FBS), L-glutamine, 100 U/mL penicillin and 100 μg/mL streptomycin (all from Thermo Fisher Scientific, USA) as described previously ^79^. All cell lines were mycoplasma-free. Fibroblasts at passage (p) 2 were used for reprogramming.

### Generation, selection and iPSC maintenance

A TECAN liquid-handling platform was used to maintain and passage iPSCs, as described in ^26^. iPSCs were generated by nucleofection with episomal vectors expressing *OCT-4*, *SOX2*, *KLF4*, *L-MYC, LIN28* and shRNA against *p53* ^80^ in feeder- and serum-free conditions using TeSR™-E7™medium as described previously ^27^. The reprogrammed cells were maintained on the automated platform using TeSR™-E7™medium, with daily medium change. Pluripotent cells were selected by sorting anti-human TRA-1-60 Microbeads ^27^. Cell number was determined; cells were subsequently plated onto vitronectin XF™-coated plates and in StemFlex™ medium. Subsequent culturing was performed on the automated platform using StemFlex™medium, which was changed every 2-3 days. Passaging was performed weekly on the automated platform using ReLeSR™ onto vitronectin XF™-coated plates as described in ^27^. TRA-1-60 quantifications were performed on a MACSQuant® Analyzer 10 immediately prior to passaging to fresh plates as described ^27^. Pluripotency was assessed by expression of the markers OCT3/4 (sc-5279, Santa Cruz Biotechnology) and TRA-1-60 (MA1-023-PE, Thermo Fisher Scientific, USA) by immunocytochemistry. Virtual karyotyping was undertaken using Illumina Human Core Exome or UK Biobank Axiom™ Arrays, as described in ^26^. The iPSC lines FSA0001, FSA0002, FSA0005, FSA0006, IST2168, IST2607, MBE1006, MBE2900, MBE2906, MBE2909, TOB0199, TOB0224, TOB0412, TOB0421, TOB0431, TOB0435, TOB0474, WAB0450, WAB0069 were characterised in ^26^.

### Differentiation of iPSCs into retinal organoids

Retinal organoids were generated following ^17^ with modifications. Briefly, on day 28 formed retinal organoids with surrounding tissue were excised using a 21G needle. They were maintained in suspension culture for 7 days in PRO medium (DMEM/F12, 1:1, L-glutamine, 1% Non-essential amino acids, Penicillin-Streptomycin 10,000 U/ml) supplemented with B27 and FGF2 (10 ng/ml). On day 35, organoids were transferred to Matrigel-coated 24-well tissue culture plates and maintained for 14 days in NDM medium (Neurobasal, 1% MEM non-essential amino acids, 1% GlutaMAX, 1% glucose (45%), Penicillin-Streptomycin 10,000 U/ml) with 2% B27 and 1% N2 added fresh. Medium was changed every 2-3 days. Optic cups were dissociated with Papain Dissociation System following the manufacturer’s instructions. Briefly, cells were harvested with papain (20 U/mL) and DNase I (2,000 U/mL) for 30 minutes at 37 °C. Subsequently, NDM medium was added at a 1:1 ratio and cells were gently triturated with a P1000 pipette followed by centrifugation (5 minutes, 300 g, 4 °C). Cells were resuspended in 1 mL of 0.1% BSA in PBS solution. Subsequently, cells were counted and assessed for viability with Trypan Blue using a Countess II automated counter, then pooled at a concentration of 1000 cells/ μL (1 × 10^6^ cells/mL). Final cell viability estimates ranged between 79-99%.

### Transcriptome profiling of single cells from retinal organoids and cell-based quality control

Multi-donor single-cell suspensions were prepared for scRNA-seq using the Chromium Single Cell 3′ Library & Gel bead kit (10x Genomics; PN-120237). Each pool was loaded onto individual wells of 10x Genomics Single Cell A Chip along with the reverse transcription (RT) master mix to generate single-cell gel beads in emulsion (GEMs). Reverse transcription was performed using a C1000 Touch Thermal Cycler with a Deep Well Reaction Module (Bio-Rad) as follows: 45 min at 53 °C; 5 min at 85 °C; hold 4 °C. cDNA was recovered and purified with DynaBeads MyOne Silane Beads (Thermo Fisher Scientific; catalog no. 37002D). Subsequently, it was amplified as follows: for 3 min at 98°C; 12× (for 15 sec at 98 °C; for 20 sec at 67 °C; for 60 sec at 72°C); for 60 sec at 72 °C; hold 4 °C followed recommended cycle number based on targeted cell number. Amplified cDNA was purified with SPRIselect beads (Beckman Coulter; catalog no. B23318) and underwent quality control following manufacturer’s instructions. Sequencing libraries for each pool were labelled with unique sample indices (SI) and combined for sequencing across two 2 × 150 cycle flow cells on an Illumina NovaSeq 6000 (NovaSeq Control Software v1.6) using S4 Reagent kit (catalog no. 20039236). Raw base calls from the sequencer then underwent demultiplexing, quality control, mapping and quantification with the Cell Ranger Single Cell Software Suite 3.1.0 by 10x Genomics (https://www.10xgenomics.com/). Processed reads from the sequencer were mapped to the *Homo sapiens* reference *hg19*/*GRCh37* from ENSEMBL (release 75), and the pipeline was run using the estimated cell count value of 20,000. scRNA-seq data from each pool underwent quality control separately in R using the *ascend* package ^69^. Cells were removed if they did not meet thresholds calculated from 3 Median Absolute Deviations (MAD) of the following statistics: total Unique Molecular Identifier (UMI) counts, number of detected genes, and fraction of mitochondrial and ribosomal transcripts to total expression.

### SNP genotype analysis and imputation

DNA was extracted from cell pellets using QIAamp DNA Mini Kit (QIAGEN, 51306) following the manufacturer’s instructions. DNA concentration was determined with SimpliNano spectrophotometer (GE Life Sciences), diluted to a final concentration of 10-15 ng/μl and genotyped on UK Biobank Axiom™ Arrays at the Ramaciotti Centre for Genomics, Sydney, Australia. Samples previously screened using Illumina arrays, were re-genotyped on UK Biobank Axiom™ Arrays. Genotype data were exported into PLINK data format using GenomeStudio PLINK Input Report Plug-in v2.1.4 and screened for SNP and individual call rates (<0.97), HWE failure (P<10^−6^), and MAF (<0.01) ^81^. Samples with excess autosomal heterozygosity or with sex-mismatch were excluded. In addition, a genetic relationship matrix from all the autosomal SNPs were generated using the GCTA tool and one of any pair of individuals with estimated relatedness larger than 0.125 were removed from the analysis ^82^. Individuals with non-European ancestry were excluded outside of an “acceptable” box of +/- 6SD from the European mean in PC1 and PC2 in a SMARTPCA analysis. The 1000G Phase 3 population was used to define the axes, and the samples were projected onto those axes. Imputation was performed on each autosomal chromosome using the Michigan Imputation Server with the Haplotype Reference Consortium panel (HRC r1.1 2016) and run using Minimac3 and Eagle v2.3 ^67,68^. Only SNPs with INFO > 0.8 were retained.

### Demultiplexing of cell pools into individual donors

*demuxlet* v1.0 ^70^ and *scrublet* v0.20 ^71^ were used to demultiplex cells from mixed-donor pools using transcriptome and genotype data. Each pool was demultiplexed separately. *demuxlet* was run with exon-only SNPs and the following arguments: “*--field GP, --geno.error = 0.01, --min-mac 1, --min-callrate 0.5, alpha = 0.5, doublet-prior = 0.5*”. Cells were initially assigned a putative donor based on the maximum likelihood of reads from scRNA-seq overlapping sets of unique variants (SNPs) mapped by genotyping*. scrublet* was then used to confirm if a cell was a neotypic doublet by building a simulation of doublets based on sampled transcriptome data and scoring the cell based on its neighbors in k-means nearest neighbor graph. A cell was designated a singlet if *scrublet* agrees, and if the posterior probability of it being a singlet in *demuxlet* is greater than 0.99. Unassigned donors, doublets and cells with ambiguous assignments were omitted from downstream analyses.

### Aggregation, normalization and dimensionality reduction of scRNA-seq datasets

The unfiltered count matrices of all batches were combined into one dataset using the *cellranger aggr* pipeline. This pipeline equalized the read depth of all batches by downsampling reads from higher-depth libraries to match the lowest depth library ^83^. Cells that had been removed from single-batch analyses due to low quality, being labeled as a doublet, or with conflicting assignments, were also removed from the combined expression matrix. The SCTransform function from Seurat (v3.0.2) was applied to the filtered count matrix to perform cell-cell and batch normalization ^84^. The fraction of mitochondrial and ribosomal expression of total expression was regressed out as a part of this step, and the top 3000 most variable genes were used to calculate residuals. The residuals were then reduced to 30 principal components (PCs) using PC Analysis (PCA). These PCs were reduced further into two dimensions, for visualization and clustering via Uniform Manifold Approximation Projection (UMAP) ^75^.

### Identification and annotation of cell subpopulations

Graph-based clustering via the Louvain algorithm that was implemented in Seurat was used to identify cell subpopulations ^85,86^. First, cell-cell Euclidean distances calculated from PCs were projected onto a K-means nearest-neighbor graph (KNN). Next, the Louvain algorithm was implemented at resolutions between 0 and 1, at 10 equal intervals of 0.1. Finally, the movement of cells between subpopulations at these resolutions were visualized on to a *clustree* plot (Figure S2A) as implemented in the *clustree* R package ^87^. Regions of stability were identified from the plot, and the resolution where this region began - 0.4, was selected as the optimal resolution. Cells were divided into 22 subpopulations at this resolution. Cells that could not be assigned to a subpopulation - singletons, were assigned to a group designated ‘Cluster 0’. To annotate each subpopulation, the combined Likelihood Ratio Test (LRT) as described by ^88^ was applied to normalized, log-transformed UMI counts. Disease-specific markers within subpopulations were identified using the Wilcoxon Rank Sum test ^89^. Genes were classified as markers if they were differentially expressed, based on the thresholds of average log_2_ fold change > |0.25| and Bonferroni-corrected p-value < 0.01.

### Identification of differentiation lineages via pseudotime analysis

Differentiation lineages were identified using the *slingshot* R package ^76^. Singletons were excluded from the trajectory, and the UMAP matrix was used as input for the ‘slingshot’ function. RPC1 was selected as the root of the trajectories due to the expression of the proliferative marker MKI67, and the end-points of trajectories were defined as the relatively mature cell subpopulations: RGC1, Cones, Interneurons, RPE and Lens.

### Identification of cis-eQTL using transcriptome and genotype data

We investigated cis-eQTL associated with each subpopulation using the R package Matrix eQTL ^90^. To prepare scRNA-seq data for this stage of analysis, the normalized, corrected UMI count matrix was split by subpopulation. The mean expression of each gene, for all cells within the subpopulation from a donor, was used to build a gene-donor count matrix. Genes that were expressed in less than 30% of the donor population were excluded, and remaining mean counts were transformed (log(x+1)). SNPs with a Minor Allele Frequency (MAF) of less than 10% were filtered from the genotype dataset. SNPs within one mega-base pairs of a gene were tested. To identify lead eQTL based on population, Matrix eQTL was run with an additive linear model using sex, age and the top six genotype principal components as covariates. To identify eQTL specific to POAG, disease status was included in the model. eQTL were determined to be significant based on a threshold of FDR < 0.05. Lead eQTLs identified using the disease model underwent additional testing using the *lm* function from R to measure the size and significance of interactions between genotype and disease status.

### Transcriptome wide association study analysis

Transcriptome-wide association study analysis (TWAS) was performed using summary statistics generated by eQTL analysis. We customized our prediction models based on the online tutorial (available through https://github.com/hakyimlab/MetaXcan) to evaluate the association between predicted gene expression and glaucoma risk using GWAS summary statistics. The gene-expression prediction models were constructed from the single-cell eQTL data using the following models: “blup”, “lasso”, “top1” and “enet”. The GWAS summary statistics from a recent multitrait meta-analysis of glaucoma were used ^8^. S-PrediXcan was used to harmonize the GWAS summary statistics and to compute the gene-level association results using each subpopulation of single-cell expression data and glaucoma GWAS summary statistics ^91^. A total of 3,573 genes (across tested subpopulations) were tested for glaucoma. To correct for multiple testing, we adjusted the gene-level association P values using the Bonferroni correction method (0.05 / (total number of tests across all subpopulations)). We then compared the transcriptome-wide association results based on scRNA-seq data to bulk RNA-seq data. The bulk retinal transcriptome data were described previously ^60^. Briefly, 406 controls and age-related macular degeneration cases that passed RNA-seq and genotyping quality control were modeled with mixed models, LASSO, or elastic net according to Gusev et al ^92^. The effect sizes from these models were used as weights to calculate the gene-trait associations.

**SUPPLEMENTAL INFORMATION TITLES AND LEGENDS**

**Figure S1. Identification and characterisation of cell subpopulations.**

**Figure S2. Variation in cell differentiation between control and POAG samples.**

**Figure S3. Pseudotime analysis across lineages and sub-lineages.**

**Figure S4. Boxplots displaying the relationships between genotype and mean gene expression for all eQTLs in RGC subpopulations with a FDR < 5** ✕ **10^−8^.**

**Table S1. Summary of single cell RNA-sequencing metrics.**

**Table S2. Cell-donor deconvolution summary metrics.**

**Table S3. Subpopulation-specific markers identified by differential expression gene analysis.**

**Table S4. Number of significant cis-eQTLs across cell subpopulations.**

**Table S5. Lead cis-eQTLs from RGC subpopulations.**

**Table S6. Statistically significant SNP-by-disease state interaction eQTLs in the RGC lineage specific subpopulations.**

**Table S7. Disease-specific differentially-expressed genes in retinal ganglion cells. Supplemental Results. Description of delineated cell subpopulations.**

