## Supplemental Information for "Retinal ganglion cell-specific genetic regulation in primary open angle glaucoma"

^#^ Equal first authors

^^^ Equal senior authors

**Figure S1. Identification and characterisation of cell subpopulations. (A)** “Clustree” representation of graph-based clustering results over range of resolutions from 0 (top row) to 1 (bottom row). Arrows indicate movement of cells from one group to another at each resolution. Minimal movement of cells between groups are indicative of grouping stability. **(B)** Feature plots of selected RPC and RGC markers across all cells. Color scale represents gene expression level in a cell.

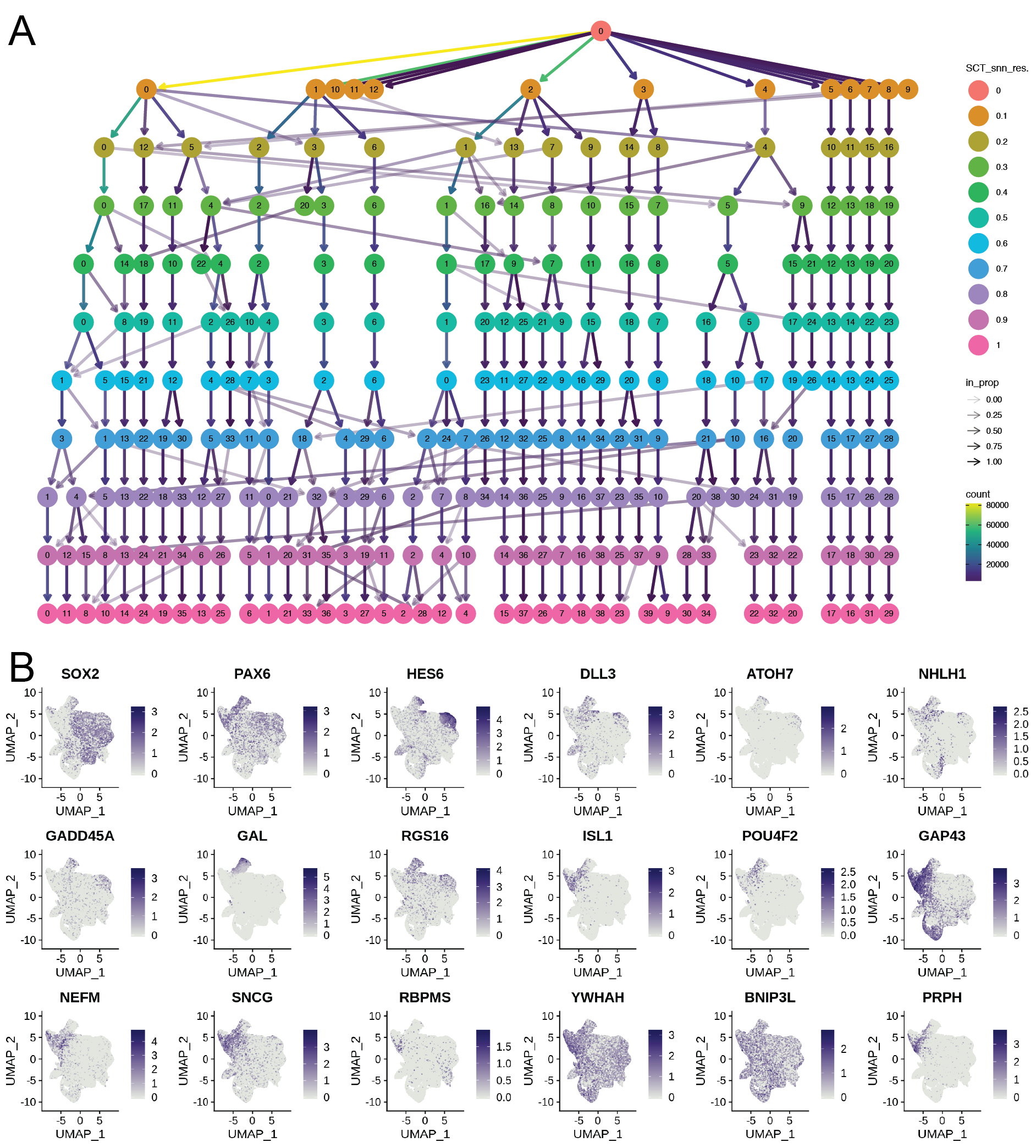

**Figure S2. Variation in cell differentiation between control and POAG samples**. (**A**) Distribution of number of cells per cell type, per donor. Each point represents a donor, with healthy donors shown in grey and donors with POAG shown in red. (**B**) Distribution of identified cell types across all donors.

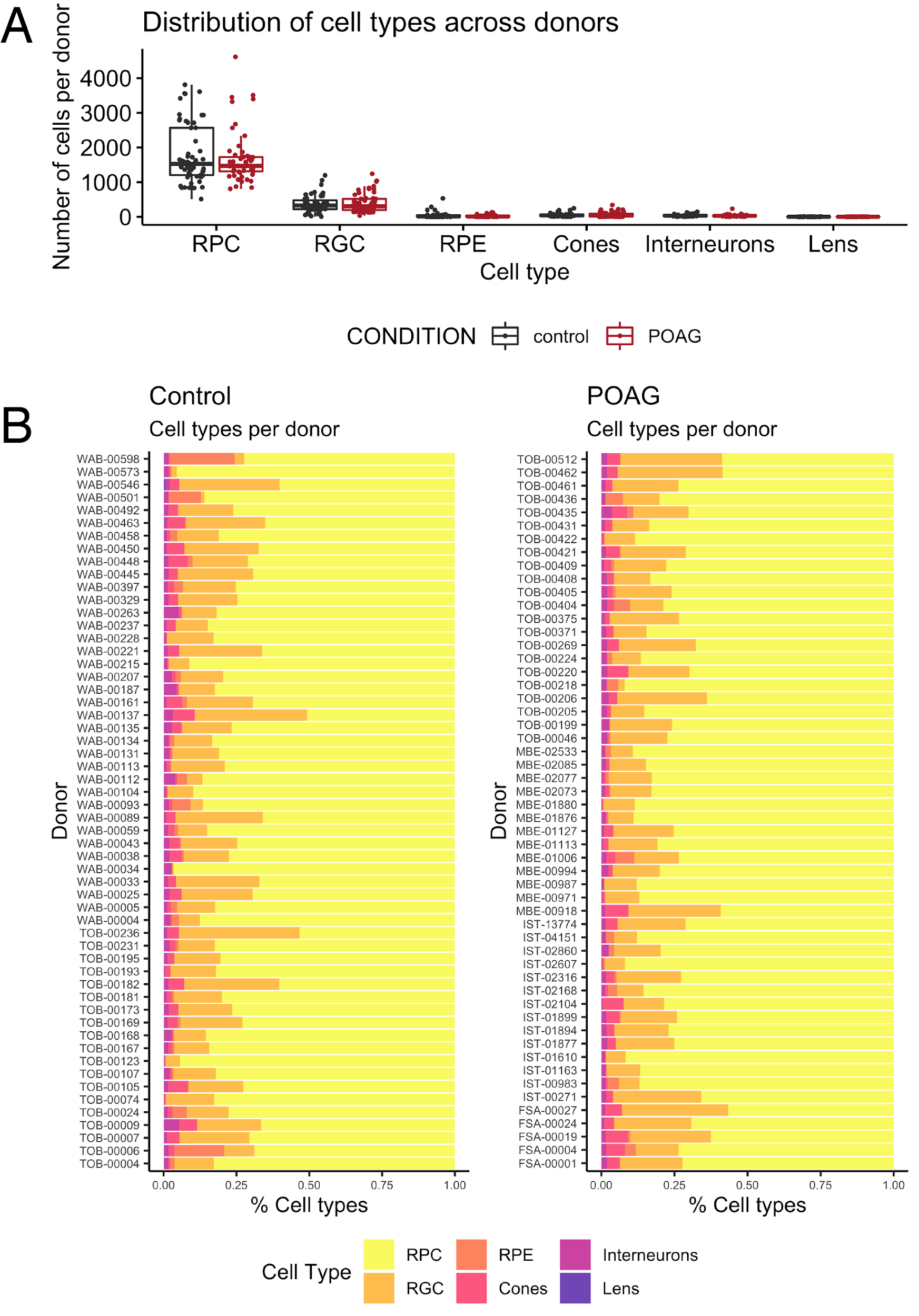

**Figure S3. Pseudotime analysis across lineages and sub-lineages.** **(A)** Lineages and sublineages of subpopulations based on relationship to RPC1, as identified by Slingshot. **(B)** Cells, colored by subpopulation are divided into sub-lineages by Slingshot and ordered based on pseudotime relative to the progenitor subpopulation, RPC1.

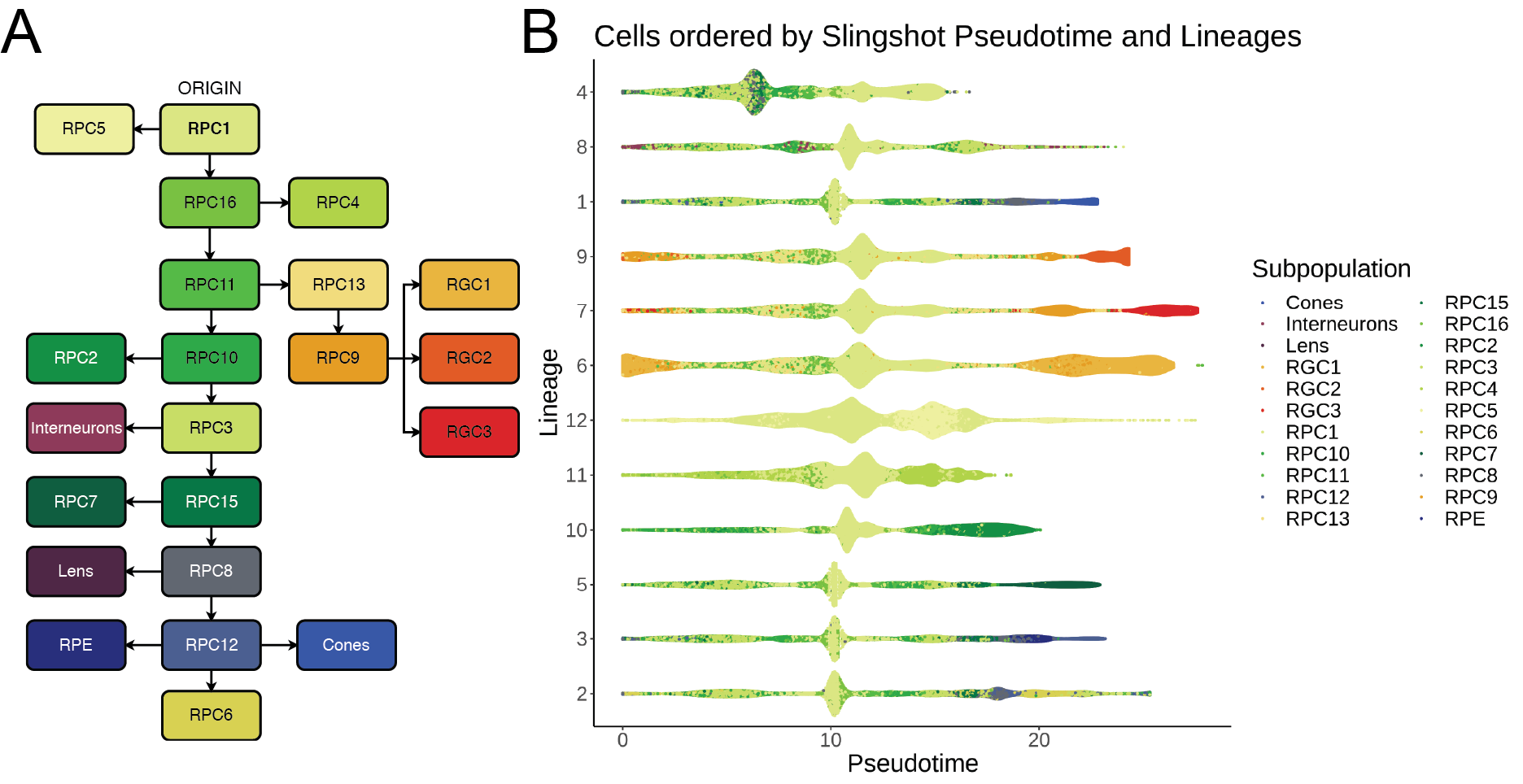

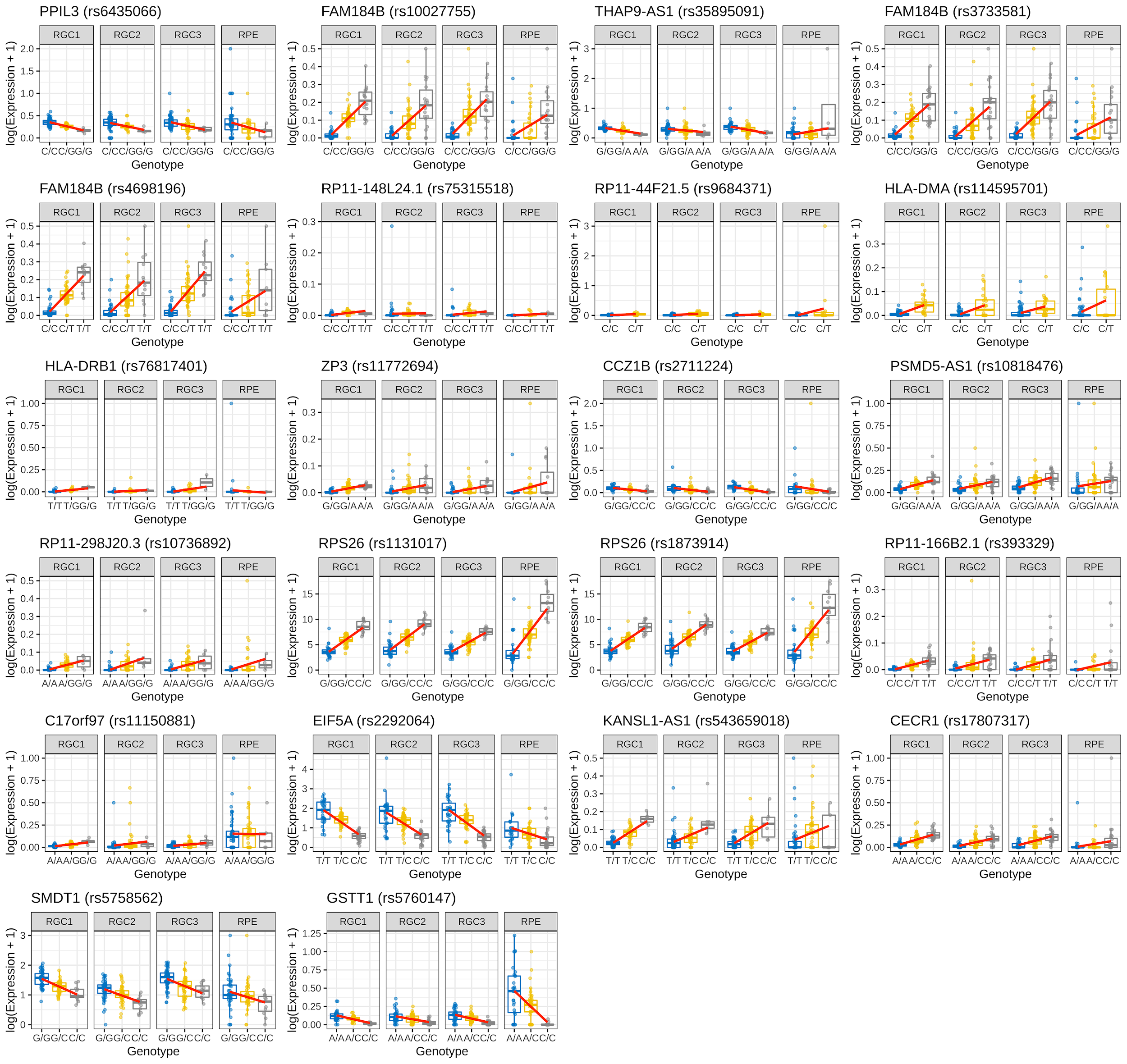

**Figure S4.** Boxplots displaying the relationships between genotype and mean gene expression for all eQTLs in RGC subpopulations with a FDR < 1 ✕ 10^-8^.

**Table S1. Summary of single cell RNA-sequencing metrics.** Single cell libraries were generated using the Chromium Single Cell Gene Expression V3 workflow by 10x Genomics. Cells derived from 160 donors were pooled and sequenced in 25 batches with an expected cell capture value of 20,000. This table summarizes the quality of read mapping and quantification of each batch.

**Table S2. Cell-donor deconvolution summary metrics.** Data from 25 multi-donor cell pools were demultiplexed with *demuxlet*. 289,617 out of 309,223 cells were traced back to donors. 29,733 cells were classified as doublets by *demuxlet* and *scrublet.* 12,786 cells were removed due to reasons described in Table S3. Pool 4 had failed as a single cell pool, and was discarded from the study.

| **Pool** | **Detected Individuals** | **Singlets** | **Doublets** |
| --- | --- | --- | --- |
| 1 | 5 | 10,630 | 1,418 |
| 2 | 7 | 15,677 | 909 |
| 3 | 6 | 10,602 | 1,027 |
| 4 | 0 | 0 | 0 |
| 5 | 5 | 10,464 | 1,107 |
| 6 | 8 | 9,652 | 485 |
| 7 | 8 | 11,106 | 716 |
| 8 | 7 | 11,586 | 1,028 |
| 9 | 7 | 12,553 | 2,280 |
| 10 | 7 | 12,725 | 1,309 |
| 11 | 7 | 10,794 | 903 |
| 12 | 7 | 9,686 | 919 |
| 13 | 8 | 13,247 | 1,135 |
| 14 | 8 | 13,662 | 1,014 |
| 15 | 6 | 13,382 | 993 |
| 16 | 8 | 13,748 | 732 |
| 17 | 8 | 10,920 | 686 |
| 18 | 7 | 14,546 | 1,314 |
| 19 | 8 | 13,504 | 508 |
| 20 | 7 | 11,402 | 786 |
| 21 | 8 | 15,621 | 856 |
| 22 | 8 | 8,883 | 546 |
| 23 | 8 | 12,306 | 728 |
| 24 | 6 | 14,648 | 1,061 |
| 25 | 3 | 8,273 | 7,273 |

**Table S3. Subpopulation-specific markers identified by differential expression gene analysis.**

**Table S4. Number of significant cis-eQTLs across cell subpopulations.**

| **Subpopulation** | **Population Model** | **Disease Model** | **Control Only** | **POAG Only** |
| --- | --- | --- | --- | --- |
| RGC1 | 297 | 280 | 117 | 106 |
| RPC1 | 456 | 440 | 115 | 155 |
| RPC2 | 136 | 139 | 100 | 88 |
| RPC3 | 143 | 148 | 34 | 65 |
| RPC4 | 144 | 136 | 77 | 63 |
| RPC5 | 222 | 217 | 82 | 68 |
| RPC6 | 69 | 63 | 36 | 21 |
| RPC7 | 44 | 50 | 49 | 44 |
| RGC2 | 45 | 41 | 10 | 29 |
| RPC8 | 66 | 73 | 26 | 17 |
| RGC3 | 89 | 95 | 33 | 23 |
| Cones | 72 | 67 | 68 | 21 |
| RPC9 | 34 | 35 | 47 | 13 |
| RPC10 | 76 | 71 | 29 | 28 |
| RPC11 | 46 | 45 | 46 | 18 |
| RPC12 | 50 | 49 | 15 | 12 |
| Interneurons | 71 | 65 | 14 | 10 |
| RPC13 | 55 | 58 | 60 | 16 |
| RPE | 48 | 38 | 39 | 30 |
| RPC14 | 24 | 25 | 13 | 28 |
| RPC15 | 47 | 40 | 15 | 12 |

**Table S5. Lead cis-eQTLs from RGC subpopulations.**

**Table S6. Statistically significant SNP-by-disease state interaction eQTLs in the RGC lineage specific subpopulations.** Abbreviation: FDR, false discovery rate.

| **Subpopulation** | **SNP** | **Gene** | **Percent expressed** | **Allelic effect** | **Standard error** | **Test statistic** | **FDR** |
| --- | --- | --- | --- | --- | --- | --- | --- |
| RGC1 | rs7222922 | *RDM1* | 0.57 | -0.022 | 0.005 | -4.626 | 1.23x10^-4^ |
| RGC1 | rs55892012 | *QSOX2* | 0.87 | -0.071 | 0.016 | -4.529 | 1.67x10^-4^ |
| RGC1 | rs4970420 | *MXRA8* | 0.93 | -0.041 | 0.009 | -4.373 | 2.68x10^-4^ |
| RGC1 | rs55892012 | *EGFL7* | 0.89 | -0.042 | 0.010 | -4.323 | 3.19x10^-4^ |
| RGC1 | rs28368130 | *CDKN2B* | 0.60 | -0.036 | 0.009 | -4.187 | 4.67x10^-4^ |
| RGC1 | rs909666 | *CYP2D6* | 0.41 | -0.005 | 0.002 | -3.423 | 4.53x10^-3^ |
| RGC1 | rs148036156 | *SGSH* | 0.86 | -0.028 | 0.008 | -3.386 | 4.92x10^-3^ |
| RGC1 | rs1475102 | *IGFBPL1* | 0.96 | 0.024 | 0.007 | 3.355 | 5.34x10^-3^ |
| RGC1 | rs449460 | *TRIM68* | 0.90 | -0.024 | 0.007 | -3.284 | 6.60x10^-3^ |
| RGC1 | rs6842644 | *C4orf27* | 0.98 | 0.088 | 0.027 | 3.264 | 6.91x10^-3^ |
| RGC1 | rs72823056 | *TUBG2* | 0.95 | -0.036 | 0.011 | -3.238 | 7.37x10^-3^ |
| RGC1 | rs7176219 | *DIS3L* | 0.97 | -0.050 | 0.015 | -3.241 | 7.37x10^-3^ |
| RGC1 | rs111945529 | *RP11-395B7.4* | 0.40 | -0.007 | 0.002 | -3.108 | 1.03x10^-2^ |
| RGC1 | rs16923133 | *NFIB* | 0.98 | -0.194 | 0.065 | -3.006 | 1.33x10^-2^ |
| RGC1 | rs13111191 | *CEP135* | 0.92 | -0.049 | 0.017 | -2.874 | 1.81x10^-2^ |
| RGC1 | rs2400439 | *RP11-373N22.3* | 0.53 | -0.007 | 0.002 | -2.837 | 1.96x10^-2^ |
| RGC1 | rs4746023 | *SAR1A* | 0.99 | 0.051 | 0.018 | 2.817 | 2.06x10^-2^ |
| RGC1 | rs12913415 | *RP11-209K10.2* | 0.34 | -0.005 | 0.002 | -2.813 | 2.07x10^-2^ |
| RGC1 | rs73507925 | *PCDH9-AS1* | 0.75 | -0.024 | 0.009 | -2.776 | 2.27x10^-2^ |
| RGC1 | rs74691737 | *NXPH1* | 0.56 | -0.026 | 0.009 | -2.771 | 2.30x10^-2^ |
| RGC1 | rs2250287 | *HLA-B* | 0.97 | -0.020 | 0.007 | -2.756 | 2.37x10^-2^ |
| RGC1 | rs17500190 | *BTBD8* | 0.76 | -0.007 | 0.003 | -2.735 | 2.47x10^-2^ |
| RGC1 | rs7776941 | *HOXA1* | 0.73 | 0.013 | 0.005 | 2.727 | 2.49x10^-2^ |
| RGC1 | rs10966858 | *RP11-125B21.2* | 0.95 | -0.012 | 0.005 | -2.611 | 3.28x10^-2^ |
| RGC1 | rs2728117 | *SPP1* | 0.98 | -0.045 | 0.017 | -2.567 | 3.64x10^-2^ |
| RGC1 | rs7756998 | *CACNG7* | 0.34 | 0.039 | 0.006 | 6.801 | 4.43x10^-8^ |
| RGC2 | rs59993224 | *TK1* | 0.76 | -0.043 | 0.014 | -2.969 | 1.19x10^-2^ |
| RGC2 | rs57656483 | *RP3-462E2.5* | 0.47 | 0.020 | 0.007 | 2.909 | 1.31x10^-2^ |
| RGC2 | rs7952972 | *PPP1R12C* | 0.53 | -0.021 | 0.007 | -2.885 | 1.37x10^-2^ |
| RGC2 | rs1985841 | *HLA-DRB1* | 0.32 | -0.032 | 0.006 | -5.294 | 8.92x10^-6^ |
| RGC2 | rs159674 | *MYO1D* | 0.33 | -0.079 | 0.023 | -3.490 | 2.62x10^-3^ |
| RGC3 | rs9268851 | *ACOX1* | 0.91 | -0.046 | 0.015 | -3.064 | 8.95x10^-3^ |
| RGC3 | rs17247560 | *RP11-298J20.3* | 0.33 | -0.017 | 0.006 | -2.869 | 1.50x10^-2^ |
| RGC3 | rs4513132 | *BAIAP2-AS1* | 0.68 | -0.055 | 0.020 | -2.734 | 2.10x10^-2^ |
| RGC3 | rs10901835 | *HPGD* | 0.49 | -0.012 | 0.005 | -2.686 | 2.35x10^-2^ |
| RGC3 | rs35278165 | *SOCS3* | 0.64 | -0.051 | 0.019 | -2.603 | 2.82x10^-2^ |
| RGC3 | rs2332691 | *RP3-525N10.2* | 0.99 | -0.045 | 0.018 | -2.526 | 3.39x10^-2^ |
| RGC3 | rs78180639 | *UVSSA* | 0.63 | -0.024 | 0.010 | -2.403 | 4.54x10^-2^ |
| RGC3 | rs12209170 | *SCPEP1* | 0.55 | -0.015 | 0.006 | -2.365 | 4.89x10^-2^ |
| RGC3 | rs11936653 | *ZNF74* | 0.80 | -0.082 | 0.022 | -3.650 | 1.50x10^-3^ |
| RGC3 | rs8070286 | *CDK12* | 0.95 | -0.098 | 0.032 | -3.031 | 7.85x10^-3^ |
| RPC13 | rs619770 | *CDK5R1* | 0.82 | 0.090 | 0.032 | 2.798 | 1.43x10^-2^ |
| RPC13 | rs11078929 | *ATG10* | 0.85 | -0.056 | 0.020 | -2.738 | 1.63x10^-2^ |
| RPC13 | rs4795749 | *FAM118A* | 0.78 | -0.060 | 0.023 | -2.676 | 1.92x10^-2^ |
| RPC13 | rs77182875 | *MNT* | 0.61 | -0.041 | 0.016 | -2.642 | 2.05x10^-2^ |
| RPC13 | rs738169 | *ARHGAP24* | 0.48 | -0.025 | 0.010 | -2.485 | 2.96x10^-2^ |
| RPC13 | rs11654664 | *AFAP1* | 0.73 | -0.092 | 0.022 | -4.262 | 1.82x10^-4^ |
| RPC13 | rs17010957 | *SPON2* | 0.33 | -0.032 | 0.010 | -3.113 | 5.62x10^-3^ |
| RPC9 | rs9291126 | *MAP3K1* | 0.53 | -0.036 | 0.013 | -2.789 | 1.35x10^-2^ |
| RPC9 | rs6827815 | *ARHGAP29* | 0.41 | -0.034 | 0.012 | -2.791 | 1.35x10^-2^ |
| RPC9 | rs13182872 | *SEZ6L* | 0.62 | 0.049 | 0.018 | 2.652 | 1.92x10^-2^ |
| RPC9 | rs12739747 | *PAPPA* | 0.53 | -0.043 | 0.016 | -2.648 | 1.92x10^-2^ |
| RPC9 | rs12170215 | *FAM227A* | 0.61 | -0.079 | 0.031 | -2.574 | 2.26x10^-2^ |
| RPC9 | rs4978658 | *FAM184B* | 0.67 | 0.041 | 0.016 | 2.506 | 2.66x10^-2^ |

**Table S7. Disease-specific differentially-expressed genes in retinal ganglion cells.**
