## Supplemental Results for "Retinal ganglion cell-specific genetic regulation in primary open angle glaucoma"

**Supplementary Results:**

**Description of Delineated Cell Clusters**

**(See Supplementary Table 2 and Figure 1A, B)**

***Subpopulation Zero*** (35,377 cells, 13.7 % of all cells, 186 conserved markers) consisted of the “orphan” samples that show no relationships at the selected resolution. Genes specific to this subpopulation, such as *SFRP2*, *DAPL1* and *CYP1B1*, *SPP1* are expressed in RPCs (Lu et al., 2019). *PLEKHA1* was also highly expressed in this subpopulation. Its expression is controlled by *PAX6* (Sun et al., 2016) which controls multipotent states of RPCs (Marquardt et al., 2001). This pattern of gene expression suggests an RPC population.

***Subpopulation One*** (29,469 cells, 13.2 % of all cells, 313 conserved markers) was characterised by genes known to be expressed in the retina. Some markers are associated with the cytoskeleton, such as *TAGLNs*, *TUBA1A*, *TUBB2A*, *ACTB*, *MYL6*, *TMSB10*. In particular, genes involved in neuronal growth (for instance *STMN2*, *GAP43*, *NEFM*) and in sensory neurons (such as *PRPH*) were enriched markers of this subpopulation. Further, some specific RGC lineage markers were also present in this cluster, including the combined expression of *STMN2/4*, *GAP43*, *PRPH*, *ISL1*, *NEFM*, *ELAVL4/HuD* and the high expression of genes enriched in RGCs (*GAP43*, *SNCG*, *POU4F1/2*, *POU6F2*, *ISL1*, *NHLH2*, *EBF1*, *EBF3*, *MYC*) (Clark et al., 2019; Freeman et al., 2011; Hu et al., 2019; Peirson et al., 2007; Rheaume et al., 2018) suggests an RGC lineage. The presence of *STMN2/*4, *PAX6*, *DAPL1, SFRP2*, *SOX2/4/11* also suggests an immature phenotype. Altogether this suggests a RGC subpopulation.

***Subpopulation Two*** (23,588 cells, 10.6 % of all cells, 142 conserved markers) was characterised by expression of *UBE2C*, *PTTG1*, *TOP2A*, *KPNA2*, *CENPF*, *HMGB2*, CCNB1 all of which participate in regulation of cell cycle (Hsiao et al., 2020; Huang et al., 2013; Loftus et al., 2017; Stros et al., 2009; Tong et al., 2007). Combined with the high expression of the proliferation marker *MKI67*, it suggests an RPC subpopulation.

***Subpopulation Three*** (20,425 cells, 9.2 % of all cells, 100 conserved markers) was characterised by genes involved in cell growth and differentiation, which, for many are not specific to the retina, such as *FABP7* (Fatty Acid Binding Protein 7, expressed in radial glial cells and immature astrocytes, as well as in retinal astrocytes and Müller cells upon injury (Chang et al., 2007)), *C1orf61*, *PTN* or *TMSB4X* (associated with cytoplasmic sequestering of NF-kappaB). The presence of the transcription factor genes *SOX2* and *SOX3* further suggests a RPC subpopulation (Taranova et al., 2006).

***Subpopulation Four*** (19,648 cells, 8.8 % of all cells, 254 conserved markers) was characterised by its main gene markers being involved in DNA binding, transcriptional activity and regulation of the cell cycle, such as *HIST1H4C*, *PTTG1*, *HMGB2*, *HMGN2*, *MKI67*, *KIAA0101/PCLAF*, *CKS1B*, *H2AFZ* or *NUSAP1*. This pattern suggests a RPC population.

***Subpopulation Five*** (15,821 cells, 7.1 % of all cells, 226 conserved markers) was characterized by expression of many ribosomal genes, including 46 *RPLs* (Ribosomal protein L), 29 *RPSs* (Ribosomal protein S) and 7 mitochondrially-encoded genes (*MT-ND3*, *MT-ND4*, *MT-CO1*, *MT-CO2*, *MT-CO3*, *MT-ATP6*, *MT-CYB*). The expression of ribosomal genes and mitochondrially-encoded genes has been correlated with development and maturation (Zhou et al., 2015). Presence of genes associated with respiratory chain suggests a metabolic switch essential for neurogenesis (Feng and Liu, 2017; Ito and Suda, 2014; Knobloch and Jessberger, 2017). This pattern of gene expression suggests a RPC population.

***Subpopulation Six*** (14,842 cells, 6.7 % of all cells) identified 288 conserved markers. Many of the most conserved markers of this subpopulation are associated with the cytoskeleton, such as *TAGLN*, *CALD1*, *TPM1*, *TPM2*, *ACTA2*, *ACTG2*, *ACTN1*, *ACTB*. Others are associated with cell differentiation and proliferation, including *LGALS1*, *CTGF/CCN2*, *ANXA2*, *S100A11*. The population expresses high levels of genes known to be expressed in early stage RPCs (including *DLX1*, *DLX2*, *ONECUT2*, *ATOH7*). This pattern is suggestive of an early RPC subpopulation.

***Subpopulation Seven*** (14,145 cells, 6.4 % of all cells, 149 conserved markers) showed varied conserved markers, with genes involved in metal-binding (*MT1X* and *MT2A*), transcription, cell cycle and proliferation (such as *HES6*, *HMGB2*, *PTTG1*, *TOP2A*). The presence of genes known to be expressed in the retina but not cell type specific as markers of this subpopulation (*CKB*), as well as markers of neuronal differentiation (*NEUROD1*) with markers of RPCs (*SFRP2*, *ASCL1*) and high levels of expression of *DLX1/2* suggests an early RPC subpopulation.

***Subpopulation Eight*** (9,744 cells, 4.4 % of all cells, 298 conserved markers) identified markers of transcriptional activity (*NFIB*, *NFIA*, *BCL11A*), and many neural markers (such as *CNTNAP2*, *LMO3*, *TUBA1A*, *TUBB2A*), involved in retinal differentiation (*SOX4*, *SOX11*), neuronal differentiation (such as *NEUROD6*, *NEUROD2*, *GPM6A*) and neuroendocrine secretion (*RTN1*). The presence of the RPC markers (*CLU, SFRP2*, *NFIA*, *NFIB* and *VIM*) together with RGC markers (*GAP43*, *PRPH*, *ELAVL4, TBR1*), combined with the high expression of genes enriched in late RPC genes (*SOX4*, *NEUROG2*), interneurons (*CALB2*), photoreceptors (*NHLH1*, *RHO*), suggest a late RPC population differentiating into retinal neurons.

***Subpopulation Nine*** (9,652 cells, 4.3 % of all cells, 355 conserved markers) showed conserved markers for nuclear transport (*UBE2C*, *KPNA2*), transcriptional activity and cell cycle (*PTTG1*, *TOP2A*, *CENPF*, *MKI67*, *CCNB1*, *CDK1*) (Hsiao et al., 2020; Huang et al., 2013; Loftus et al., 2017; Tong et al., 2007; Yu, 2007). It was also characterised by the RPC/RGC marker *ATOH7* and RGC gene markers *GAP43*, *PRPH*, *NEFM* and *ELAVL4*. Combined with the high expression of RPC genes (*DLX1/2*, *GAL*, *ONECUT2*, *ATOH7*), this suggests an RPC population differentiating into a RGC population.

***Subpopulation Ten*** (7,843 cells, 3.5 % of all cells, 202 conserved markers) shows expression of markers of neurogenesis including of the retina (*NEUROD1*/*4*, *C8orf46/VXN*), transcription (*PRDM1*), melanogenesis (*DCT*), apoptosis (*PHLDA1*), which suggests a progenitor population (Lu et al., 2019). The presence of conserved markers that are RPC markers together with photoreceptor progenitor markers (such as *OTX2* and *CRX*), and with the low expression of genes associated with most cell types of the retina suggests a differentiating RPC population.

***Subpopulation 11*** (7,693 cells, 3.5 % of all cells, 96 conserved markers) shows conserved marker genes associated with RPCs (*CLU*, *SFRP2, VIM*, *DLX2* and *SOX4* (Lu et al., 2019)) as well as conserved markers associated with neuronal development (*NNAT*, *MEG3*, *PEG10*, *MEIS2*), neuronal differentiation and growth (*SOX4*, *MLLT11*, *STMN2*, NSG2), neuroendocrine secretion (*RTN1, PCSK1N*), cytoskeleton organization (*TUBA1A*, *STMN1*, *TUBB2A/B*, *MARCKSL*1, *DCX*, TMSB10), telomere maintenance (*TERF2IP*), transport (*VAMP2*). We observed high expression of genes enriched in amacrine cells (ONECUT1/2*, ESRRB*) also known to be expressed by RGCs (Rheaume et al., 2018; Sapkota et al., 2014). Together with the high expression of genes enriched in RGCs (*GAP43*, *SNCG*, *POU4F1, POU4F2,* *POU6F2*, *ISL1*, *NHLH2*, , *EBF1/3*, *MYC*), this pattern of expression suggests a RGC population.

***Subpopulation 12*** (7,455 cells, 3.3 % of all cells, 42 conserved markers) identified conserved markers associated with hormonal activity (*TTR*, *IGFBP7*, *C1orf194*), calcium signaling (*TRPM3*), ciliogenesis (*FAM183A*, *C1orf192/ CFAP126*), cell adhesion/ Wnt signalling (*TPBG*, *PIFO*), retinal development (RP11-356K23.1) or cytoskeleton organization (*TPPP3*). Together with the high expression of genes found in photoreceptor progenitors (*OTX2*, *CRX*) and cone cells (*LHX9*, *ARR3*, *GNGT2*, *GUCA1C*, *DCT*, *LMO4*, *THRB*, *RXRG*), this pattern suggests a cone population.

***Subpopulation 13*** (7,321 cells, 3.3 % of all cells, 121 conserved markers) identified conserved markers associated with neuronal development (*HOXB4/5/6*, *TAGLN3*), transcriptional regulation (*RP11-834C11.4*, *HOTAIRM1*, *HOXB-AS3*), neuronal growth (*NEFM*, STMN2), synaptic transmission (*SNCG*, *LAMP*5), axon guidance (*NOVA1*), cytoskeletal organization (*TUBB2B, TUBA1A*). Expression of *HOX* genes suggests an RPC population and expression of genes associated with RGCs, (*TAGLN3*, *NEFM*, *STMN2*, *SNCG)* (Hu et al., 2019; Laboissonniere et al., 2017, 2019) suggests a RPC population differentiating into RGCs.

***Subpopulation 14*** (6,962 cells, 3.1 % of all cells, 347 conserved markers) identified genes associated with synaptic transmission (*SST*, *NSG1*, *SNCA/SNCG*), neuronal growth (*STMN2*, *GAP43*, *NEFM*, *ISL1*, *PRPH*, *ELAVL4/HuD*), neuroendocrine secretion (*RTN1*) which are also markers of RGCs. Yet the low level of expression of cell type specific markers suggests a RPC population.

***Subpopulation 15*** (6,399 cells, 2.9 % of all cells, 137 conserved markers) was characterized by expression of several genes involved in lipid metabolism and trafficking, i.e. *APOA1*, *APOA2*, *APOC1*, *APOC3*, *APOE*, *NPC2*. It was also suggested that apolipoproteins may be important for membrane assembly during cell division (Grehan et al., 2001). We also detected genes associated with regulation of cell cycle: *KRT8/18* (Toivola et al., 2001), *S100A10* (Wang et al., 2012), *FTL* (Ferritin light chain) (Bogdan et al., 2016), *AFP* (Mizejewski, 2016), *KRT19* (Sharma et al., 2019) and maintenance of open chromatin - *HMGA1* (Ozturk et al., 2014). This pattern of gene expression suggests a RPC population.

***Subpopulation 16*** (4,429 cells, 2.0 % of all cells, 100 conserved markers) identified genes involved in neuromodulation (*NTS*), neuronal differentiation (*NEUROD6*, *NFIB*, *NFIA*, *CALB2*, *BCL11A*, *GPM6A*, *NEUROD2*, *NSG2*, *GAP43*, *TBR1*), neuroendocrine secretion (*RTN1*), transport (*FXYD6*, *LY6H*, *VAMP2*), synaptic transmission (*CAMKV*, *GRIA2*, MEF2C), axonal outgrowth/ autophagy (*FEZ1*), cell adhesion (*CNTNAP2*), and cytoskeleton organization (*THSD7A*, *DSTN*]). This indicates a RPC population.

***Subpopulation 17*** (4,012 cells, 1.8 % of all cells) only identified 34 conserved markers, with some associated with differentiation (*IGFBP5*, *FABP7*, *HES6, STMN2*). The high expression of *TFAP2A/B*, *CALB1* and *CHAT* suggests an interneuron population, such as horizontal and amacrine cells (Kaewkhaw et al., 2015).

***Subpopulation 18*** (3,590 cells, 1.6 % of all cells) showed 32 conserved markers genes, including markers associated with neuronal differentiation (*HES6, FABP7, ANXA2, PCP4, CALB1*), signalling (*CXCL14*, *TTYH1,* *TPBG, SFRP2*), metabolism (*GATM, CKB, HMGCS1*), or transcriptional activity (*HMGB2, ID2, HMGN2, CKS2, PTTG1, CENPF, NUSAP1, CDK1*), cytoskeleton organization (*TMSB4X, TUBA1B, STMN4*). The high expression of *VSX2*  further suggests a RPC population (Zou and Levine, 2012).

***Subpopulation 19*** (3,326 cells, 1.5 % of all cells, 635 conserved markers) identified genes associated with RPE cells, such as *TTR*, *TRPM3*, *IGFBP7*, *CST3,* *RPE65*, *RBP1/CRBP1*, or *SERPINF1/PEDF*. It is also characterised by genes involved in early retinal development, including the RPE and eye morphogenesis (*SOX4*, *SOX11*, *BMP7*, *GJA1*, *PTN*). Many RPE genes are highly expressed in this population. Altogether, this suggests a RPE population.

***Subpopulation 20*** (2,936 cells, 1.3 % of all cells, 128 conserved markers) identified genes associated with protein transport such as *CRYAB, HSPA5* and *HSPB1*; iron homeostasis (*FTL*), gene regulation (*NEAT1*) or cytoskeleton regulation (such as *STMN1, TUBB* or *ACTB*). The absence of markers for specific cell types suggests a RPC population.

***Subpopulation 21*** (2,866 cells, 1.3 % of all cells, 129 conserved markers) identified genes associated with early neural differentiation (*HES4*, *HES6, CLU , PAX6, POU4F2, RORB, DLX1/2*, *SOX11*), axon guidance (*CXCR4*), neurite growth (*MDK, RTN4*), synapse formation (*NPTX2*), synaptic plasticity (*SERPINI1*), synaptic vesicle transport (*SLC18A2, CPLX2*) and neuromodulation (*GAL*, *TRH*). The expression of early RPE genes suggests a RPC population with potential for neuronal and RPE differentiations.

***Subpopulation 22*** (528 cells, 0.2 % of all cells, 33 conserved markers) is characterised by 12 genes encoding different types of crystallins as well as *LIM2* which is highly expressed in the lens. The presence of *AQP5* and *MIP*, *CYP26A1* all known to play roles in the lens further supports the lens identity of this subpopulation.

Lidgerwood, G.E., Senabouth, A., Smith-Anttila, C.J.A., Gnanasambandapillai, V., Kaczorowski, D.C., Amann-Zalcenstein, D., Fletcher, E.L., Naik, S.H., Hewitt, A.W., Powell, J., et al. Evolving Transcriptomic Signature of Human Pluripotent Stem Cell-Derived Retinal Pigment Epithelium Cells With Age.

Ligon, K.L., Huillard, E., Mehta, S., Kesari, S., Liu, H., Alberta, J.A., Bachoo, R.M., Kane, M., Louis, D.N., Depinho, R.A., et al. (2007). Olig2-regulated lineage-restricted pathway controls replication competence in neural stem cells and malignant glioma. Neuron *53*, 503–517.

Lin, M., Pedrosa, E., Shah, A., Hrabovsky, A., Maqbool, S., Zheng, D., and Lachman, H.M. (2011). RNA-Seq of human neurons derived from iPS cells reveals candidate long non-coding RNAs involved in neurogenesis and neuropsychiatric disorders. PLoS One *6*, e23356.

Liu, W., and Hajjar, K.A. (2016). The annexin A2 system and angiogenesis. Biol. Chem. *397*, 1005–1016.

Liu, Y., Myrvang, H.K., and Dekker, L.V. (2015). Annexin A2 complexes with S100 proteins: structure, function and pharmacological manipulation. Br. J. Pharmacol. *172*, 1664–1676.

Loftus, K.M., Cui, H., Coutavas, E., King, D.S., Ceravolo, A., Pereiras, D., and Solmaz, S.R. (2017). Mechanism for G2 phase-specific nuclear export of the kinetochore protein CENP-F. Cell Cycle *16*, 1414–1429.

Lou, Y., Jiang, H., Cui, Z., Wang, X., Wang, L., and Han, Y. (2018). Gene microarray analysis of lncRNA and mRNA expression profiles in patients with high‑grade ovarian serous cancer. International Journal of Molecular Medicine.

Lu, Y., Shiau, F., Yi, W., Lu, S., Wu, Q., Pearson, J.D., Kallman, A., Zhong, S., Hoang, T., Zuo, Z., et al. (2019). Single-cell analysis of human retina identifies evolutionarily conserved and species-specific mechanisms controlling development (bioRxiv).

Lujan, E., Chanda, S., Ahlenius, H., Südhof, T.C., and Wernig, M. (2012). Direct conversion of mouse fibroblasts to self-renewing, tripotent neural precursor cells. Proc. Natl. Acad. Sci. U. S. A. *109*, 2527–2532.

Mahoney, S.-A., Hosking, R., Farrant, S., Holmes, F.E., Jacoby, A.S., Shine, J., Iismaa, T.P., Scott, M.K., Schmidt, R., and Wynick, D. (2003). The Second Galanin Receptor GalR2 Plays a Key Role in Neurite Outgrowth from Adult Sensory Neurons. The Journal of Neuroscience *23*, 416–421.

Mao, X., An, Q., Xi, H., Yang, X.-J., Zhang, X., Yuan, S., Wang, J., Hu, Y., Liu, Q., and Fan, G. (2019). Single-Cell RNA Sequencing of hESC-Derived 3D Retinal Organoids Reveals Novel Genes Regulating RPC Commitment in Early Human Retinogenesis. Stem Cell Reports *13*, 747–760.

Marquardt, T., Ashery-Padan, R., Andrejewski, N., Scardigli, R., Guillemot, F., and Gruss, P. (2001). Pax6 is required for the multipotent state of retinal progenitor cells. Cell *105*, 43–55.

Martínez, P., and Blasco, M.A. (2011). Telomeric and extra-telomeric roles for telomerase and the telomere-binding proteins. Nat. Rev. Cancer *11*, 161–176.

Martinsson-Ahlzén, H.-S., Liberal, V., Grünenfelder, B., Chaves, S.R., Spruck, C.H., and Reed, S.I. (2008). Cyclin-dependent kinase-associated proteins Cks1 and Cks2 are essential during early embryogenesis and for cell cycle progression in somatic cells. Mol. Cell. Biol. *28*, 5698–5709.

de Melo, J., Du, G., Fonseca, M., Gillespie, L.-A., Turk, W.J., Rubenstein, J.L.R., and Eisenstat, D.D. (2005). Dlx1 and Dlx2 function is necessary for terminal differentiation and survival of late-born retinal ganglion cells in the developing mouse retina. Development *132*, 311–322.

Miller, D.J., and Fort, P.E. (2018). Heat Shock Proteins Regulatory Role in Neurodevelopment. Front. Neurosci. *12*, 821.

Mizejewski, G.J. (2016). The alpha-fetoprotein (AFP) third domain: a search for AFP interaction sites of cell cycle proteins. Tumour Biol. *37*, 12697–12711.

Mongiat, M., Andreuzzi, E., Tarticchio, G., and Paulitti, A. (2016). Extracellular Matrix, a Hard Player in Angiogenesis. Int. J. Mol. Sci. *17*.

Moore, K.B., Logan, M.A., Aldiri, I., Roberts, J.M., Steele, M., and Vetter, M.L. (2018). C8orf46 homolog encodes a novel protein Vexin that is required for neurogenesis in Xenopus laevis. Dev. Biol. *437*, 27–40.

Morcillo, J., Martínez-Morales, J.R., Trousse, F., Fermin, Y., Sowden, J.C., and Bovolenta, P. (2006). Proper patterning of the optic fissure requires the sequential activity of BMP7 and SHH. Development *133*, 3179–3190.

Morishima, K., Matsuura, S., Tauchi, H., Nakamura, A., and Komatsu, K. (1999). A polymorphic CA repeat marker at the human 27-kD calbindin (CALB1) locus. J. Hum. Genet. *44*, 414–415.

Mu, X., Fu, X., Sun, H., Beremand, P.D., Thomas, T.L., and Klein, W.H. (2005). A gene network downstream of transcription factor Math5 regulates retinal progenitor cell competence and ganglion cell fate. Dev. Biol. *280*, 467–481.

Mu, X., Fu, X., Beremand, P.D., Thomas, T.L., and Klein, W.H. (2008). Gene regulation logic in retinal ganglion cell development: Isl1 defines a critical branch distinct from but overlapping with Pou4f2. Proc. Natl. Acad. Sci. U. S. A. *105*, 6942–6947.

Nelson, B.R., Hartman, B.H., Georgi, S.A., Lan, M.S., and Reh, T.A. (2007). Transient inactivation of Notch signaling synchronizes differentiation of neural progenitor cells. Dev. Biol. *304*, 479–498.

Orosz, F., and Ovádi, J. (2008). TPPP orthologs are ciliary proteins. FEBS Lett. *582*, 3757–3764.

Ozturk, N., Singh, I., Mehta, A., Braun, T., and Barreto, G. (2014). HMGA proteins as modulators of chromatin structure during transcriptional activation. Front Cell Dev Biol *2*, 5.

Pan, L., Deng, M., Xie, X., and Gan, L. (2008). ISL1 and BRN3B co-regulate the differentiation of murine retinal ganglion cells. Development *135*, 1981–1990.

Paraoan, L., Grierson, I., and Maden, B.E. (2000). Analysis of expressed sequence tags of retinal pigment epithelium: cystatin C is an abundant transcript. Int. J. Biochem. Cell Biol. *32*, 417–426.

Passmore, L.A., Schmeing, T.M., Maag, D., Applefield, D.J., Acker, M.G., Algire, M.A., Lorsch, J.R., and Ramakrishnan, V. (2007). The eukaryotic translation initiation factors eIF1 and eIF1A induce an open conformation of the 40S ribosome. Mol. Cell *26*, 41–50.

Peirson, S.N., Oster, H., Jones, S.L., Leitges, M., Hankins, M.W., and Foster, R.G. (2007). Microarray analysis and functional genomics identify novel components of melanopsin signaling. Curr. Biol. *17*, 1363–1372.

Peng, Y.-R., Shekhar, K., Yan, W., Herrmann, D., Sappington, A., Bryman, G.S., van Zyl, T., Do, M.T.H., Regev, A., and Sanes, J.R. (2019). Molecular Classification and Comparative Taxonomics of Foveal and Peripheral Cells in Primate Retina. Cell *176*, 1222–1237.e22.

Pirondi, S., Fernandez, M., Schmidt, R., Hökfelt, T., Giardino, L., and Calzà, L. (2005). The galanin-R2 agonist AR-M1896 reduces glutamate toxicity in primary neural hippocampal cells. J. Neurochem. *95*, 821–833.

Poliak, S., Salomon, D., Elhanany, H., Sabanay, H., Kiernan, B., Pevny, L., Stewart, C.L., Xu, X., Chiu, S.-Y., Shrager, P., et al. (2003). Juxtaparanodal clustering of Shaker-like K+ channels in myelinated axons depends on Caspr2 and TAG-1. J. Cell Biol. *162*, 1149–1160.

Radeke, M.J., Radeke, C.M., Shih, Y.-H., Hu, J., Bok, D., Johnson, L.V., and Coffey, P.J. (2015). Restoration of mesenchymal retinal pigmented epithelial cells by TGFβ pathway inhibitors: implications for age-related macular degeneration. Genome Med. *7*, 58.

Ramírez-Castillejo, C., Sánchez-Sánchez, F., Andreu-Agulló, C., Ferrón, S.R., Daniel Aroca-Aguilar, J., Sánchez, P., Mira, H., Escribano, J., and Fariñas, I. (2006). Pigment epithelium–derived factor is a niche signal for neural stem cell renewal. Nature Neuroscience *9*, 331–339.

Raz, A., and Goodman, D.S. (1969). The interaction of thyroxine with human plasma prealbumin and with the prealbumin-retinol-binding protein complex. J. Biol. Chem. *244*, 3230–3237.

Rheaume, B.A., Jereen, A., Bolisetty, M., Sajid, M.S., Yang, Y., Renna, K., Sun, L., Robson, P., and Trakhtenberg, E.F. (2018). Single cell transcriptome profiling of retinal ganglion cells identifies cellular subtypes. Nat. Commun. *9*, 2759.

Rivera, L.B., and Brekken, R.A. (2011). SPARC promotes pericyte recruitment via inhibition of endoglin-dependent TGF-β1 activity. J. Cell Biol. *193*, 1305–1319.

Rodgers, H.M., Belcastro, M., Sokolov, M., and Mathers, P.H. (2016). Embryonic markers of cone differentiation. Mol. Vis. *22*, 1455–1467.

Rodríguez, P., Higueras, M.A., González-Rajal, A., Alfranca, A., Fierro-Fernández, M., García-Fernández, R.A., Ruiz-Hidalgo, M.J., Monsalve, M., Rodríguez-Pascual, F., Redondo, J.M., et al. (2012). The non-canonical NOTCH ligand DLK1 exhibits a novel vascular role as a strong inhibitor of angiogenesis. Cardiovasc. Res. *93*, 232–241.

Rouillard, A.D., Gundersen, G.W., Fernandez, N.F., Wang, Z., Monteiro, C.D., McDermott, M.G., and Ma’ayan, A. (2016). The harmonizome: a collection of processed datasets gathered to serve and mine knowledge about genes and proteins. Database *2016*.

Saito, Y., Miranda-Rottmann, S., Ruggiu, M., Park, C.Y., Fak, J.J., Zhong, R., Duncan, J.S., Fabella, B.A., Junge, H.J., Chen, Z., et al. (2016). NOVA2-mediated RNA regulation is required for axonal pathfinding during development. Elife *5*.

Samuel, W., Kutty, R.K., Vijayasarathy, C., Pascual, I., Duncan, T., and Redmond, T.M. (2010). Decreased expression of insulin-like growth factor binding protein-5 during N-(4-hydroxyphenyl)retinamide-induced neuronal differentiation of ARPE-19 human retinal pigment epithelial cells: regulation by CCAAT/enhancer-binding protein. J. Cell. Physiol. *224*, 827–836.

Sapkota, D., Chintala, H., Wu, F., Fliesler, S.J., Hu, Z., and Mu, X. (2014). Onecut1 and Onecut2 redundantly regulate early retinal cell fates during development. Proc. Natl. Acad. Sci. U. S. A. *111*, E4086–E4095.

Schmitt, S., Aftab, U., Jiang, C., Redenti, S., Klassen, H., Miljan, E., Sinden, J., and Young, M. (2009). Molecular Characterization of Human Retinal Progenitor Cells. Investigative Opthalmology & Visual Science *50*, 5901.

Sharma, P., Alsharif, S., Bursch, K., Parvathaneni, S., Anastasakis, D.G., Chahine, J., Fallatah, A., Nicolas, K., Sharma, S., Hafner, M., et al. (2019). Keratin 19 regulates cell cycle pathway and sensitivity of breast cancer cells to CDK inhibitors. Sci. Rep. *9*, 14650.

Shekhar, K., Lapan, S.W., Whitney, I.E., Tran, N.M., Macosko, E.Z., Kowalczyk, M., Adiconis, X., Levin, J.Z., Nemesh, J., Goldman, M., et al. (2016). Comprehensive Classification of Retinal Bipolar Neurons by Single-Cell Transcriptomics. Cell *166*, 1308–1323.e30.

Sindhu Kumari, S., Gupta, N., Shiels, A., FitzGerald, P.G., Menon, A.G., Mathias, R.T., and Varadaraj, K. (2015). Role of Aquaporin 0 in lens biomechanics. Biochem. Biophys. Res. Commun. *462*, 339–345.

Sleat, D.E., Wiseman, J.A., El-Banna, M., Price, S.M., Verot, L., Shen, M.M., Tint, G.S., Vanier, M.T., Walkley, S.U., and Lobel, P. (2004). Genetic evidence for nonredundant functional cooperativity between NPC1 and NPC2 in lipid transport. Proc. Natl. Acad. Sci. U. S. A. *101*, 5886–5891.

Sridhar, A., Hoshino, A., Finkbeiner, C.R., Chitsazan, A., Dai, L., Haugan, A.K., Eschenbacher, K.M., Jackson, D.L., Trapnell, C., Bermingham-McDonogh, O., et al. (2020). Single-Cell Transcriptomic Comparison of Human Fetal Retina, hPSC-Derived Retinal Organoids, and Long-Term Retinal Cultures. Cell Rep. *30*, 1644–1659.e4.

Storch, J., and Xu, Z. (2009). Niemann–Pick C2 (NPC2) and intracellular cholesterol trafficking. Biochimica et Biophysica Acta (BBA) - Molecular and Cell Biology of Lipids *1791*, 671–678.

Stros, M., Polanska, E., Struncova, S., and Pospisilova, S. (2009). HMGB1 and HMGB2 proteins up-regulate cellular expression of human topoisomerase II. Nucleic Acids Research *37*, 2070–2086.

Strunnikova, N.V., Maminishkis, A., Barb, J.J., Wang, F., Zhi, C., Sergeev, Y., Chen, W., Edwards, A.O., Stambolian, D., Abecasis, G., et al. (2010). Transcriptome analysis and molecular signature of human retinal pigment epithelium. Hum. Mol. Genet. *19*, 2468–2486.

Sun, J., Zhao, Y., McGreal, R., Cohen-Tayar, Y., Rockowitz, S., Wilczek, C., Ashery-Padan, R., Shechter, D., Zheng, D., and Cvekl, A. (2016). Pax6 associates with H3K4-specific histone methyltransferases Mll1, Mll2, and Set1a and regulates H3K4 methylation at promoters and enhancers. Epigenetics Chromatin *9*, 37.

Suryadinata, R., Sadowski, M., Steel, R., and Sarcevic, B. (2011). Cyclin-dependent kinase-mediated phosphorylation of RBP1 and pRb promotes their dissociation to mediate release of the SAP30·mSin3·HDAC transcriptional repressor complex. J. Biol. Chem. *286*, 5108–5118.

Szuchet, S., Nielsen, J.A., Lovas, G., Domowicz, M.S., de Velasco, J.M., Maric, D., and Hudson, L.D. (2011). The genetic signature of perineuronal oligodendrocytes reveals their unique phenotype. Eur. J. Neurosci. *34*, 1906–1922.

Tanegashima, K., Suzuki, K., Nakayama, Y., Tsuji, K., Shigenaga, A., Otaka, A., and Hara, T. (2013). CXCL14 is a natural inhibitor of the CXCL12-CXCR4 signaling axis. FEBS Lett. *587*, 1731–1735.

Taranova, O.V., Magness, S.T., Fagan, B.M., Wu, Y., Surzenko, N., Hutton, S.R., and Pevny, L.H. (2006). SOX2 is a dose-dependent regulator of retinal neural progenitor competence. Genes Dev. *20*, 1187–1202.

Tasic, B., Yao, Z., Graybuck, L.T., Smith, K.A., Nguyen, T.N., Bertagnolli, D., Goldy, J., Garren, E., Economo, M.N., Viswanathan, S., et al. (2018). Shared and distinct transcriptomic cell types across neocortical areas. Nature *563*, 72–78.

Thijssen, V.L., and Griffioen, A.W. (2014). Galectin-1 and -9 in angiogenesis: a sweet couple. Glycobiology *24*, 915–920.

Thomas, A.G., and Henry, J.J. (2014). Retinoic acid regulation by CYP26 in vertebrate lens regeneration. Dev. Biol. *386*, 291–301.

Thompson, A., Berry, M., Logan, A., and Ahmed, Z. (2019). Activation of the BMP4/Smad1 Pathway Promotes Retinal Ganglion Cell Survival and Axon Regeneration. Invest. Ophthalmol. Vis. Sci. *60*, 1748–1759.

Thorenoor, N., Faltejskova-Vychytilova, P., Hombach, S., Mlcochova, J., Kretz, M., Svoboda, M., and Slaby, O. (2016). Long non-coding RNA ZFAS1 interacts with CDK1 and is involved in p53-dependent cell cycle control and apoptosis in colorectal cancer. Oncotarget *7*, 622–637.

Toivola, D.M., Nieminen, M.I., Hesse, M., He, T., Baribault, H., Magin, T.M., Omary, M.B., and Eriksson, J.E. (2001). Disturbances in hepatic cell-cycle regulation in mice with assembly-deficient keratins 8/18. Hepatology *34*, 1174–1183.

Tong, Y., Tan, Y., Zhou, C., and Melmed, S. (2007). Pituitary tumor transforming gene interacts with Sp1 to modulate G1/S cell phase transition. Oncogene *26*, 5596–5605.

Trimarchi, J.M., Stadler, M.B., and Cepko, C.L. (2008). Individual retinal progenitor cells display extensive heterogeneity of gene expression. PLoS One *3*, e1588.

Tsigelny, I.F., Kouznetsova, V.L., Lian, N., and Kesari, S. (2016). Molecular mechanisms of OLIG2 transcription factor in brain cancer. Oncotarget *7*, 53074–53101.

Turano, C., Coppari, S., Altieri, F., and Ferraro, A. (2002). Proteins of the PDI family: unpredicted non-ER locations and functions. J. Cell. Physiol. *193*, 154–163.

Ujike, H., Takaki, M., Kodama, M., and Kuroda, S. (2002). Gene expression related to synaptogenesis, neuritogenesis, and MAP kinase in behavioral sensitization to psychostimulants. Ann. N. Y. Acad. Sci. *965*, 55–67.

Urade, Y., and Hayaishi, O. (2000). Biochemical, structural, genetic, physiological, and pathophysiological features of lipocalin-type prostaglandin D synthase. Biochim. Biophys. Acta *1482*, 259–271.

Vanlandewijck, M., He, L., Mäe, M.A., Andrae, J., Ando, K., Del Gaudio, F., Nahar, K., Lebouvier, T., Laviña, B., Gouveia, L., et al. (2018). Author Correction: A molecular atlas of cell types and zonation in the brain vasculature. Nature *560*, E3.

Wallimann, T., Wyss, M., Brdiczka, D., Nicolay, K., and Eppenberger, H.M. (1992). Intracellular compartmentation, structure and function of creatine kinase isoenzymes in tissues with high and fluctuating energy demands: the “phosphocreatine circuit” for cellular energy homeostasis. Biochem. J *281 ( Pt 1)*, 21–40.

Wang, C.-Y., Chen, C.-L., Tseng, Y.-L., Fang, Y.-T., Lin, Y.-S., Su, W.-C., Chen, C.-C., Chang, K.-C., Wang, Y.-C., and Lin, C.-F. (2012). Annexin A2 silencing induces G2 arrest of non-small cell lung cancer cells through p53-dependent and -independent mechanisms. J. Biol. Chem. *287*, 32512–32524.

Wang, G., Huang, W., Li, W., Chen, S., Chen, W., Zhou, Y., Peng, P., and Gu, W. (2018a). TFPI-2 suppresses breast cancer cell proliferation and invasion through regulation of ERK signaling and interaction with actinin-4 and myosin-9. Sci. Rep. *8*, 14402.

Wang, P., Li, J., Zhao, W., Shang, C., Jiang, X., Wang, Y., Zhou, B., Bao, F., and Qiao, H. (2018b). A Novel LncRNA-miRNA-mRNA Triple Network Identifies LncRNA RP11-363E7.4 as An Important Regulator of miRNA and Gene Expression in Gastric Cancer. Cell. Physiol. Biochem. *47*, 1025–1041.

Wang, Y., Dang, Y., Liu, J., and Ouyang, X. (2016). The function of homeobox genes and lncRNAs in cancer. Oncol. Lett. *12*, 1635–1641.

Wilson, S.W., and Houart, C. (2004). Early Steps in the Development of the Forebrain. Developmental Cell *6*, 167–181.

Winkler, E.A., Bell, R.D., and Zlokovic, B.V. (2011). Central nervous system pericytes in health and disease. Nature Neuroscience *14*, 1398–1405.

Yan, W., Laboulaye, M.A., Tran, N.M., Whitney, I.E., Benhar, I., and Sanes, J.R. Molecular identification of sixty-three amacrine cell types completes a mouse retinal cell atlas.

Yang, Z., Ding, K., Pan, L., Deng, M., and Gan, L. (2003). Math5 determines the competence state of retinal ganglion cell progenitors. Dev. Biol. *264*, 240–254.

Yoshida, K., Kawamura, K., and Imaki, J. (1993). Differential expression of c-fos mRNA in rat retinal cells: regulation by light/dark cycle. Neuron *10*, 1049–1054.

Yu, H. (2007). Cdc20: a WD40 activator for a cell cycle degradation machine. Mol. Cell *27*, 3–16.

Yu, R.T., Chiang, M.Y., Tanabe, T., Kobayashi, M., Yasuda, K., Evans, R.M., and Umesono, K. (2000). The orphan nuclear receptor Tlx regulates Pax2 and is essential for vision. Proc. Natl. Acad. Sci. U. S. A. *97*, 2621–2625.

Yuniati, L., Scheijen, B., van der Meer, L.T., and van Leeuwen, F.N. (2019). Tumor suppressors BTG1 and BTG2: Beyond growth control. J. Cell. Physiol. *234*, 5379–5389.

Zagozewski, J.L., Zhang, Q., Pinto, V.I., Wigle, J.T., and Eisenstat, D.D. (2014). The role of homeobox genes in retinal development and disease. Dev. Biol. *393*, 195–208.

Zhao, P.Y., Gan, G., Peng, S., Wang, S.-B., Chen, B., Adelman, R.A., and Rizzolo, L.J. (2015). TRP Channels Localize to Subdomains of the Apical Plasma Membrane in Human Fetal Retinal Pigment Epithelium. Invest. Ophthalmol. Vis. Sci. *56*, 1916–1923.

Zhou, X., Liao, W.-J., Liao, J.-M., Liao, P., and Lu, H. (2015). Ribosomal proteins: functions beyond the ribosome. J. Mol. Cell Biol. *7*, 92–104.

Zou, C., and Levine, E.M. (2012). Vsx2 controls eye organogenesis and retinal progenitor identity via homeodomain and non-homeodomain residues required for high affinity DNA binding. PLoS Genet. *8*, e1002924.
